# Early-life social experience shapes social avoidance reactions in larval zebrafish

**DOI:** 10.1101/2020.03.02.972612

**Authors:** Antonia H. Groneberg, João C. Marques, A. Lucas Martins, Gonzalo G. de Polavieja, Michael B. Orger

## Abstract

Social experiences greatly define successive social behavior. Lack of such experiences, especially during critical phases of development, can severely impede the ability to behave adequately in social contexts. To date it is not well characterized how early-life social isolation leads to social deficits and impacts development. In many model species, it is challenging to fully control social experiences, because they depend on parental care. Moreover, complex social behaviors involve multiple sensory modalities, contexts, and actions. Hence, when studying social isolation effects, it is particularly important to parse apart social deficits from general developmental effects, such as abnormal motor learning. Here, we characterized how social experiences during early development of zebrafish larvae modulate their social behavior, at one week of age, when social avoidance reactions can be measured as discrete swim events. We show that raising larvae in social isolation leads to enhanced social avoidance, in terms of reaction distance and reaction strength. Specifically, larvae raised in isolation use a high-acceleration escape swim bout, the short latency C-start, more frequently during social interactions. These behavioral differences are absent in non-social contexts. By ablating the lateral line and presenting the fish with local water vibrations, we show that lateral line inputs are both necessary and sufficient to drive enhanced social avoidance reactions. Taken together, our results show that social experience during development is a critical factor in shaping mechanosensory avoidance reactions in larval zebrafish.

**Highlights:**

- Larval zebrafish raised in isolation show enhanced social avoidance reactions
- Enhanced avoidance is composed of increased avoidance distances and usage of high acceleration escape swims
- The lateral line sensory organ is necessary and sufficient for the increased usage of high acceleration escape swims

## Introduction

An important aspect of behavior is that it is adaptive and can change according to previous outcomes of the actions of the self or others. During early development, as the nervous system builds and refines its connections, an animal is particularly sensitive to external sensory input, or lack thereof (e.g. reviewed by Andersen [2003]). Social experiences during early development can have long lasting effects on social and other behaviors, as shown by the devastating phenotype of rhesus monkeys raised with social deprivation described by Harlow et al. [1965]. Since then, numerous studies have reported effects of social isolation raising in a variety of species, including rodents [Heidbreder et al., 2000, Lukkes et al., 2009], cockroaches [Lihoreau et al., 2009], lizards [Ballen et al., 2014], mites [Schausberger et al., 2017] and fish [Hesse et al., 2015, Shams et al., 2018]. Yet, the underlying mechanisms of how social experiences shape behavior and brain development have yet to be fully understood. One reason for this is a lack of full control over social experiences in species with parental care, where social isolation along with maternal separation can interfere with regular development. Moreover, in most animals, social behavior is complex, being triggered by multiple interacting sensory cues [Chen and Hong, 2018] and leading to behavioral outputs which constitute composite actions that can be difficult to isolate [Anderson and Perona, 2014]. Thus, the resulting phenotype of developmental social isolation is composed of various symptoms, making social deficits hard to differentiate from general impairments in motor development.

Zebrafish are a social species, aggregating in groups in the wild [Engeszer et al., 2007, Spence and Smith, 2007] and in the laboratory [Arganda et al., 2012, Dreosti et al., 2015, Stednitz et al., 2018]. They develop oviparously, without parental care, making them an ideal vertebrate species to control precisely the social experiences during development. An early study from the 1940s reported that adult zebrafish show social attraction even without prior social experience [Breder and Halpern, 1946]. Similarly, juvenile zebrafish raised in social isolation show regular levels of visual attraction to a projected, naturalistically moving dot [Larsch and Baier, 2018]. This suggests that social attraction develops without the need for social experiences. However, it has also been shown that zebrafish prefer familiar over unfamiliar visual social cues [Engeszer et al., 2004] and that olfactory kin preference requires visual and olfactory social experiences [Hinz et al., 2013]. Moreover, group cohesion in adult zebrafish was shown to be lower after social isolation-raising [Shams et al., 2018]. Therefore, unlike the drive for social attraction, fine-tuned aspects of social behavior seems to be acquired through social experiences.

Apart from social attraction, moving in social groups also requires keeping the right distance to neighbors. This has been formalized as a rule of avoidance, according to which an individual aims at maintaining a minimum distance towards others [Couzin et al., 2002, Inada and Kawachi, 2002]. In developing zebrafish, social attraction develops gradually over the course of the second and third week of development [Hinz and de Polavieja, 2017, Dreosti et al., 2015, Larsch and Baier, 2018, Stednitz and Washbourne, 2020]. Social avoidance, on the other hand, can be observed robustly in one-week-old larvae [Hinz and de Polavieja, 2017, Marques et al., 2018, Mirat et al., 2013].

Despite having a simplified social behavior, there are advantages to studying one-week-old zebrafish larvae. First, at this age fish are amenable to non-invasive manipulations, including bath application of psychoactive drugs [Rihel et al., 2010], genetic and optical ablation of defined sets of neurons [Asakawa and Kawakami, 2008, Sternberg et al., 2016], and optogenetic control of neural activity [Douglass et al., 2008, Friedrich et al., 2010]. Second, larvae are small and transparent enough to enable whole brain imaging with single neuron resolution in behaving animals [Feierstein et al., 2015], including freely moving fish [Kim et al., 2017, Marques et al., 2020, Cong et al., 2017], an ideal situation where animals can interact with each other. Finally, larval zebrafish organize their behaviors, including social interactions, in sequences of discrete bouts [Budick and O’Malley, 2000, Fero et al., 2011], which can be detected automatically and classified into types [Mirat et al., 2013, Marques et al., 2018, Mearns et al., 2020, Johnson et al., 2020], enabling precise behavioral phenotyping.

Here, we characterized how the social experience during early development affects social avoidance behavior in one-week-old larval zebrafish. Using high-speed video tracking and unsupervised clustering of swim bout types in freely interacting larvae [Marques et al., 2018], we found that isolation-raised larvae avoid conspecifics at larger distances by executing short latency C-starts [Burgess and Granato, 2007a] with higher probability. These behavioral differences are absent in non-social contexts, including upon the presentation of visual and acoustic stimuli that elicit escape reactions. Ablation of the lateral line reduces enhanced avoidance reactions, and the presentation of local water vibrations, which mimic the movements of another fish, elicits them. Therefore, lateral line inputs are both necessary and sufficient to drive enhanced social avoidance reactions. In summary, our results show that lack of social experience during development affects the type of avoidance response that fish execute, and is therefore a critical factor in shaping mechanosensory avoidance reactions in larval zebrafish.

## Results

### Isolation raised larval zebrafish show enhanced social avoidance reactions

To measure social interactions in one-week-old larvae, we tracked the positions and tail movements of groups of seven wild-type zebrafish larvae swimming freely in a circular arena under constant, homogeneous illumination (Fig. 1 A). Testing larvae were either raised in groups (GR) or in social isolation from 0 days post fertilization (dpf) until testing (ISO). We computed the density distribution of positions of all other larvae relative to a focal larva and compared it to a position density of a randomized distribution. In line with previous reports [Hinz and de Polavieja, 2017, Marques et al., 2018] there is a low density region centered around the focal larva where it is less likely to find other larvae swimming (Fig. 1 B). This region has been termed ‘avoidance area’ and appears to be larger across the ISO groups than the GR groups. In order to quantify the avoidance area for each replicate per recording group, we detected the contour of the central area as defined by a common cutoff in density ratio and computed its area (Fig. 1 B right panel). Thereby we found that ISO groups have a larger avoidance area than GR fish (Fig. 1 C; *p <* 0.0001, N=9).

**Figure 1:**
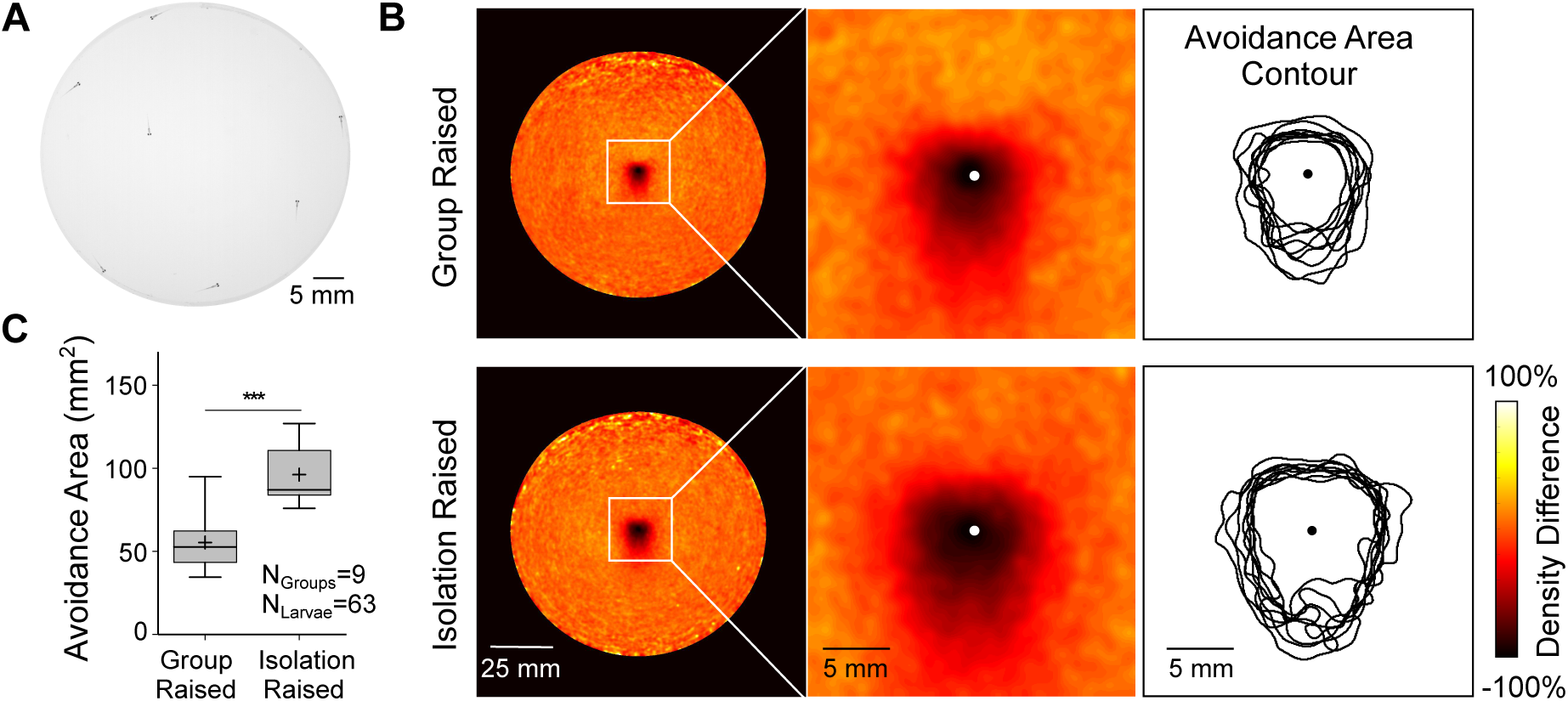
Isolation-raised larvae have a larger social avoidance area. (A) Groups of seven larvae raised either in groups or in isolation were video recorded for 1 h while swimming in a circular arena 50 mm in diameter. (B) Density distribution between real and temporally shuffled data for group-raised (top row) and isolation-raised larvae (bottom row). The center panel shows a zoom in section of the density distribution. The dot in the center indicates the position of the focal fish facing upwards. The avoidance area was quantified as the area where the data point density distribution is below 60% of the shuffled data. Shown on the left are the contours of the avoidance area calculated for each replicate of groups of seven larvae overlaid per raising condition. (C) Box plots depict the 25-75-percentile of avoidance area measures comparing the effect of the raising condition. Whiskers signal the range of data and + indicates the mean per condition. Sample size is shown in the inset and asterisks signal the results of a two-tailed unsigned Wilcoxon test between raising conditions (alpha level 0.05).

In order to study in greater detail how larvae react to one another, we adjusted the behavioral assay to test pairs of larvae. We used a recording chamber with four separate swimming arenas (Fig. S1 A). In line with the results from the group recordings, the low density region of neighboring positions in pairs of ISO larvae is larger than in GR larvae (Fig. S1 B). Because the density distribution for pairs of larvae was more sparse, we computed a one dimensional measure of avoidance by subtracting the distance between the two fish in a pair (henceforth referred to as the inter-fish distance) from a random distribution (see Methods for detail). Thereby we quantified that, also when tested in pairs, ISO fish show a larger avoidance distance than GR fish (Fig. S1 C, *p* = 0.0069, *N* = 43, see Table S1).

Locomotion in larval zebrafish is composed of discontinuous bouts of sequential tail deflections, which can be classified in 13 kinematically different swim bout types [Marques et al., 2018]. We tracked the tail movements (Fig. S1 D) of freely swimming pairs of larvae and applied this bout classification based on 73 kinematic parameters per swim bout, as described previously [Marques et al., 2018]. For simplicity, we show here only two of the kinematic parameters extracted from the tail movements that define a swim bout (Fig. 2 A). The C1 angle measures the change in heading caused by the first tail beat, while the displacement sums over the distance swum by the larva when integrating the path of the bout. Based on these two parameters, we can distinguish between three major classes of movements. Forward displacing bout types, such as slow1, slow2, the short capture swim (short CS) and the approach swim (AS) have a low C1 angle and variable, but relatively low levels of displacement. Reorienting bout types, such as the J-turn, high angle turn (HAT) and routine turn (RT) also show a relatively low displacement, but combined with a larger C1 angle than forward displacing bouts. And, lastly, there are six bout types that displace the larvae one body length (*∼*4.2 mm) or more and have variable C1 angles. This last set of bouts comprises the long capture swim (long CS), burst swim (BS), the shadow avoidance turn (SAT), the O-bend and the long and short latency C-starts (LLC and SLC, respectively).

**Figure 2.**
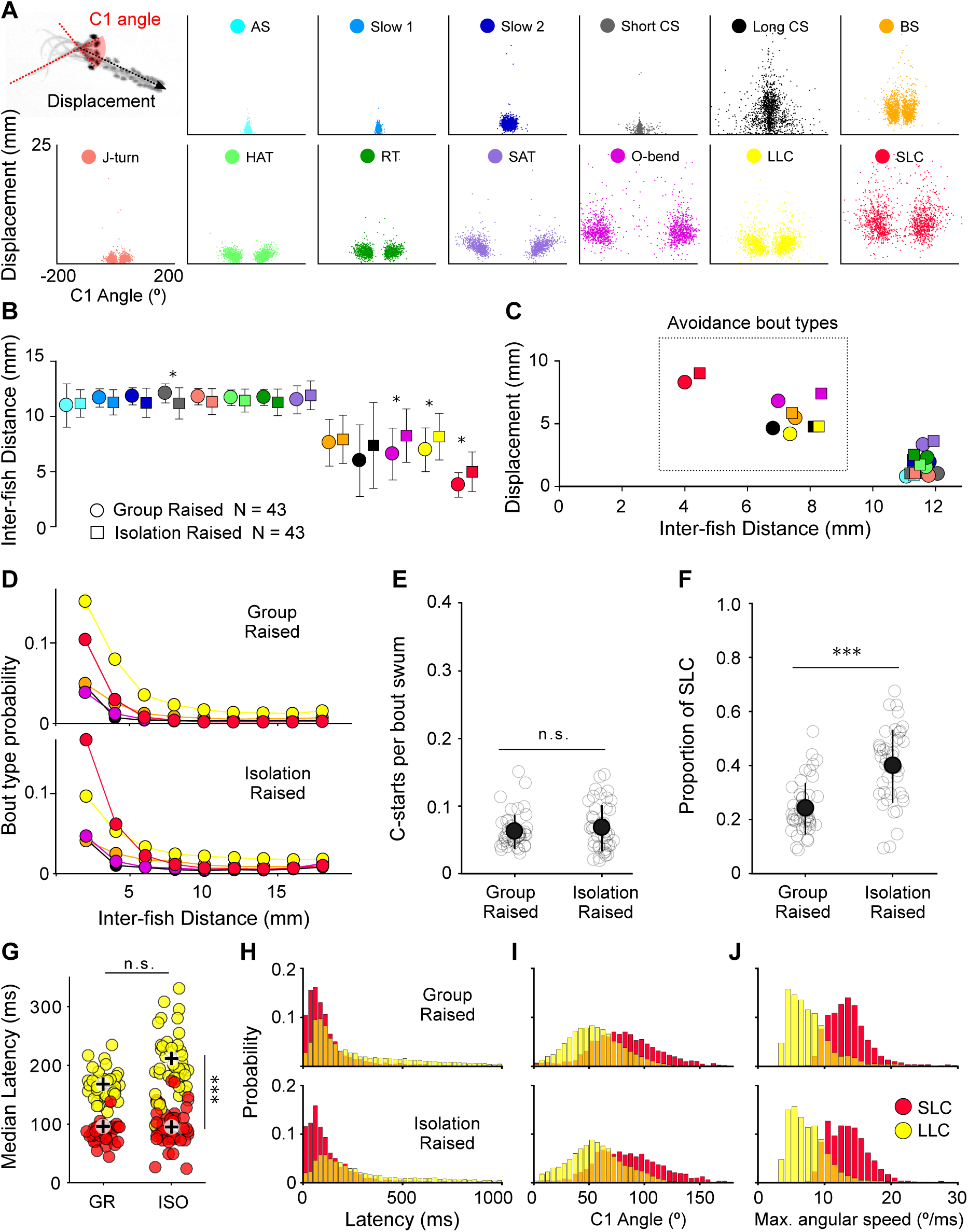
Isolation raised larvae perform large displacement bouts at larger distance and use more SLCs. (A) Overview of the 13 swim bout types of larval zebrafish. C1 angle measures the change in heading caused by the first half beat of the tail relative to the axis prior to the bout. Displacement measures the sum over the distance swum by the larva by integrating the path of the bout. Shown are scatter plots of 1500 bouts per bout type category randomly chosen from 43 pairs of group-raised larvae. (B) The median distance between the two larvae per pair of each bout type is compared between group and isolation-raised pairs. Bout types are color-coded, markers and errorbars show mean standard deviation, for group raised (circles) and isolation raised larvae (squares). Asterisks signal significant results of Bonferroni-Holm corrected p-values from unsigned Wlicoxon tests between raising conditions for each bout type (alpha level 0.05). (C) Displacement per bout type is plotted against the mean inter-fish distance, revealing that large displacing bout types tend to occur at small inter-fish distances. (D) Probability of usage of the close-range bout types (O-bend, BS, long CS, LLC and SLC) is expressed in proportion of each bout type over all 13 bout types. This proportion is plotted for inter-fish distance bins of 2mm. Shown are mean values across 43 pairs of larvae per raising condition. (E) The number of C-starts is calculated as the sum over LLC and SLC, normalized to the total number of bouts performed by each larval pair. (F) The proportion of SLC of all C-starts as measured per testing pair for each raising condition. (E,F) Light gray open circles show every replicate per condition, dark gray filled circles and error-bars signal mean standard deviation. Asterisk specify the results of unsigned Wlicoxon tests between raising conditions. (G) Latency was calculated as the time difference between the C-start onset and the onset of the previous bout performed by the non-focal larva. Shown are median latency measures per larval pair for group raised (GR) and isolation raised larvae (ISO), color-coded as red for SLC and yellow for LLC. Asterisks indicate the summarized results of unsigned Wilcoxon tests between raising conditions (horizontal) and signed Wilcoxon tests between C-start types (vertical), after Bonferroni-Holm correction for multiple comparison; alpha level 0.05. (H-J) Histograms of latency, C1 angle and maximum angular speed per C-start type, as pooled over animals per raising conditions; group raised top row; isolation raised bottom row. Red bars show probability values for SLCs, while yellow bars depict LLC values. AS: Approach Swim; Long CS: Long capture swim; BS: Burst swim; HAT: High angle turn; RT: Routine turn; SAT: Shadow avoidance turn; LLC: Long latency C-start; SLC: Short latency C-start.

To understand if and how all 13 bout types are being used during social interactions, we calculated the median inter-fish distance per bout type for each pair of fish. While the majority of bout types show a median value similar to the radius of the area, five bout types are shifted towards smaller inter-fish distances, suggesting that they are used during close-range social interactions. Multiple comparison of raising condition for each bout type revealed that three of these close-range bout types (O-bend, LLC and SLC) are performed at increased inter-fish distances by ISO larvae (Fig. 2 B, see Table S2). Interestingly, the close-range bout types show the largest displacement (Fig. 2 C), suggesting that larvae use those large displacing bouts to avoid one another.

We next compared the probability of using these close-range bouts as a function of interfish distance (Fig. 2 D). As expected there is an inverse relationship between bout type usage and inter-fish distance. Yet, there was a striking difference between the two raising conditions. While both show a preference for C-starts over the other close-range bout types, the relative choice of C-start bout type differs. We found that GR larvae use more LLCs (*p* = 0.0003), while ISO larvae are more likely to use SLCs (*p* = 0.0005). The usage of long CS, BS and O-bend were not different between raising conditions (see Table S3 for full statistical comparison).

To test if ISO fish are generally more reactive to a social stimulus, we compared the overall number of C-starts (sum over LLC and SLC), normalized to the total number of bouts swum. There was no difference between raising conditions (Fig. 2 E, *p* = 0.28, see Table S1), but specifically the proportion of SLCs of all C-starts was increased in ISO larvae (Fig. 2 F, *p <* 0.0001, see Table S1).

Long and short latency C-starts have originally been described in response acoustic startles [Burgess and Granato, 2007a]. As evident from their respective tail angle traces, first two tail beats occur with a larger acceleration in SLCs than LLCs (Fig. S1 D). Thus, the SLC can be considered a high-acceleration version of the C-start. Moreover, as suggested by their names, the two C-starts occur at a different latency after a sudden acoustic stimulus. We therefore tested if we can also find a difference in latency in the socially induced C-starts. Because there is no clearly defined stimulus onset for socially triggered C-starts, we measured the time delay between the onset of the C-start of the focal larvae and the last preceding bout of the non-focal larva. The frequency distribution of these measures of reaction latency are not as clearly separated as expected for acoustic startles. Nonetheless, the median latency per larval pair was higher for LLCs than for SLCs in both raising conditions (see Table S4), hence matching their attributed names of long and short latency bouts (Fig. 2 G-H). Note that there was no significant difference in latency between GR and ISO, suggesting that ISO larvae react similarly, but at a larger inter-fish distance (see Fig. 2 B) and with a farther displacing bout type.

In order to confirm that the two C-start types produced by GR and ISO larvae belong to equivalent bout types, we mapped the bouts in the original principal component space of kinematic parameters that was used for the classification (Fig. S1 E). LLCs and SLCs of both raising conditions were overlapping in the principal component space, showing that they were classified correctly and that GR and ISO C-starts do not represent different bout types. To validate that the bout types are similar to the originally described acoustic startle C-starts, we next compared the C1 angle and the angular speed. Along with the latency, these two measures were originally used to differentiate LLCs and SLCs [Burgess and Granato, 2007a]. For both measures we found a significant effect of C-start bout type, but not of raising condition (Fig. 2 I-J, see Table S4). These results indicate that the bout types themselves are not different between GR and ISO larvae, but rather only their preferred usage in a social context.

### Isolation-raising affects general locomotion

Next, we tested for general effects of isolation raising on locomotion. For this we compared GR and ISO larvae swimming alone or in pairs in the same testing arena as described above. The overall distribution of bout type choices in pairs was similar between GR and ISO larvae (Fig. S2 A top panel), with the exception of the SAT, BS, O-bend, LLC and SLC (see Table S5). However, these differences disappear when testing the larvae alone in the recording arena (Fig. S2 A bottom panel, see Table S5). These results suggest that ISO larvae use more large displacement bouts specifically in social contexts. Independent of the social testing context, ISO larvae swim fewer and longer bouts (Fig. S2 B-E, see Table S6). We observed similar overall locomotion effects when treating GR larvae after exposure to the generic dopamine agonist, apomorphin (Fig. S2 F-G, see Table S6). This treatment, however, had no effect on the social avoidance measures (Fig. S2 H-I, see Table S7), suggesting that general locomotion and social avoidance effects are dissociated. Indeed, we found that isolating larvae only from 3dpf until testing at 6dpf reduced general locomotion effects (Fig. S2 J-K, see Table S6), while maintaining the social avoidance phenotype (Fig. S2 L-M, see Table S7). For subsequent experiments we applied this adjusted protocol of social isolation from 3dpf onward.

### Vision and mechanosensation contribute differently to social avoidance reactions

As an entry point to the mechanism of social avoidance reactions, we analyzed the angle of the non-focal larva position upon C-start onset (Fig. 3 A, see Table S8). The angular distribution was skewed towards the tail direction of the focal larva, in the blind spot of the larval vision. Fish can sense proximate water motion through their mechanosensory lateral line (LL), composed of neuromasts distributed laterally along the body axis and around the head [Bleckmann and Zelick, 2009]. We next aimed at testing the relative contribution of vision and LL sensing to social avoidance reactions and the isolation phenotyope. First, we tested larval pairs in darkness, where, as expected, the proportion of C-starts with the nonfocal fish positioned behind the tail increased for both GR and ISO larvae (Fig. 3 D, see Table S8). The avoidance area was reduced when testing in darkness and compared to light, in line with previous reports [Marques et al., 2018]. Furthermore, we no longer observed an increased avoidance area in ISO larvae (Fig. 3 E, *p* = 0.235), while the difference in SLC proportion between raising conditions persisted (Fig. 3 F, *p <* 0.0001; see Table S9). These results suggest that LL mechanosensation contributes to the choice of C-start bout type. We therefore next ablated the LL using neomycin, an ototoxic agent that was shown to ablate the hair cells of the LL, but not the inner ear [Buck et al., 2012]. In line with reduced LL sensing, the angular distribution of non-focal fish position upon C-start onsets was no longer skewed towards the tail direction after neomycin treatment (Fig. 3 G, see Table S8). The avoidance area was of similar magnitude as in control fish, but the difference between raising conditions was diminished (Fig. 3 H, *p* = 0.281). Also the difference in SLC proportion between GR and ISO was reduced in magnitude, albeit still present after neomycin treatment (Fig. 3 I, *p* = 0.003, see Table S9). To validate these findings, we used a second ablation reagent, *CuSO*_4_, which yielded similar results (Fig. S3 A-C), while a control incubation with regular fish medium maintained the observed social avoidance phenotype of ISO larvae (Fig. S3 D-F). To summarize these results, we calculated the effect size of the difference between GR and ISO testing animals per treatment (Fig. 3 J). We hypothesise that the escape distance during social avoidance is informed by input from the visual and the LL system, while the choice of C-start escape bout type only depends on LL mechanosenation (Fig. 3 K).

**Figure 3:**
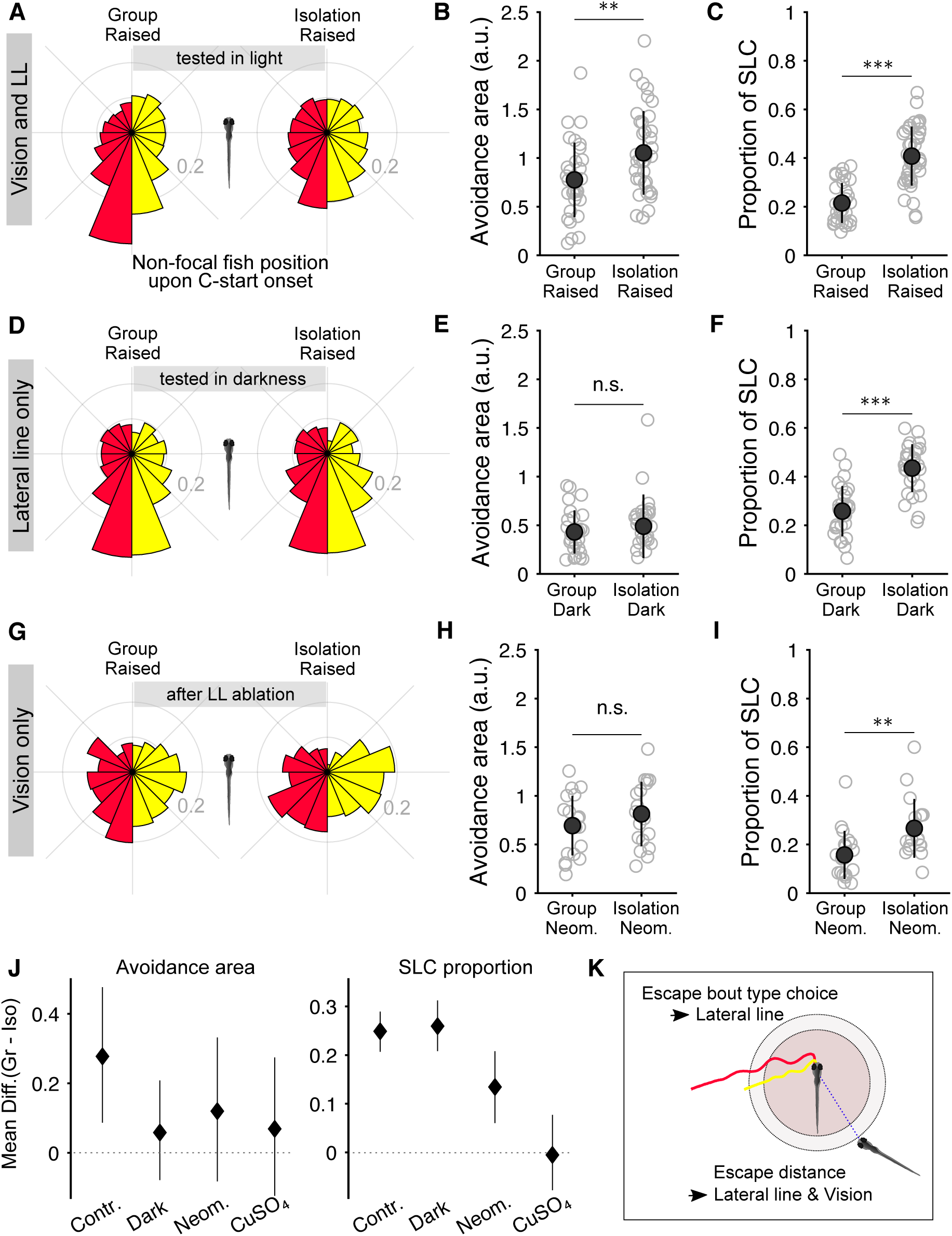
Contribution of vision and lateral line senses to enhanced social avoidance reactions in isolation-raised larvae. (A-C) Pairs of larvae raised in groups or in isolation were tested while swimming freely with homogeneous background illumination; *N*_GR_ = 31; *N*_ISO_ = 39. (D-F) Larval pairs were tested in complete darkness; *N*_GR_ = 27; *N*_ISO_ = 28. (G-I) Larvae were incubated for 1h in neomycin and given a 2 hour recovery period prior to testing in pairs with homogeneous background illumination; *N*_GR_ = 19; *N*_ISO_ = 18. (A,D,E) Circular histograms of the position of the non-focal larva upon the onset of the focal C-start. Data is pooled over fish per raising condition and treatment. Red and yellow bars show data for SLC and LLC, respectively. (B,E,H) Social avoidance area as calculated from the distribution of distances between the two fish throughout the recording period. (C,F,I) The proportion of SLC bouts over the sum of SLC and LLC. Light gray open circles show each replicate per condition, dark gray filled circles and error-bars signal mean standard deviation. Asterisks specify the results of unsigned Wilcoxon tests between raising conditions, alpha level 0.05. (J) Effect sizes of the difference between group and isolation raised larvae, calculated as the mean difference calculated with a bootstrapping method. (K) Summary of the proposed contribution of vision and lateral line mechanosensation to the social avoidance phenotype of isolation-raised larvae.

### Controlled water vibrations reproduce social avoidance reactions

To further decipher the contribution of vision and LL sensing to social avoidance reactions and the ISO avoidance phenotype, we next tested behavioural responses to controlled, single sensory-modality stimuli presented in closed-loop to larvae swimming alone in the testing arena. First, we presented radially expanding dark shadows, which have previously been shown to produce escape responses [Temizer et al., 2015, Dunn et al., 2016, Bhattacharyya et al., 2017, Marques et al., 2018]. Such stimuli were effective at triggering O-bend and SAT bout type, but not C-starts in GR (Fig. 4 A), as well as ISO fish (Fig. S4 A). The response probability of these two bout types was similar for GR and ISO fish (Fig. 4 B, *p* = 0.43) and there was no difference in the relative bout choice between raising conditions (Fig. 4 C, *p* = 0.9; see Table S10). We then tested a dark spot of constant size approaching the larva at a steady speed of 0.5 cm/sec. Unlike the expanding stimuli, this approaching dot protocol proved to be highly effective in triggering SLCs in GR (Fig. 4 D) and ISO fish (Fig. S4 B). We found that ISO fish have a lower overall response rate of C-starts with the approaching dot (Fig. 4 E, *p* = 0.0003), but that when they respond they do so with a similar proportion of SLC bout types as GR fish (Fig. 4 F, *p* = 0.24). Furthermore, when comparing the C-start latency, equivalent to a measure of reaction distance from the dot, GR and ISO fish were similar (Fig. S4 D, *p* = 0.77 and *p* = 0.78 for SLC and LLC, see Table S11). These results are in line with our hypothesis that a visual stimulus alone is insufficient to produce the avoidance distance phenotype and does not account for the different C-start choice.

**Figure 4:**
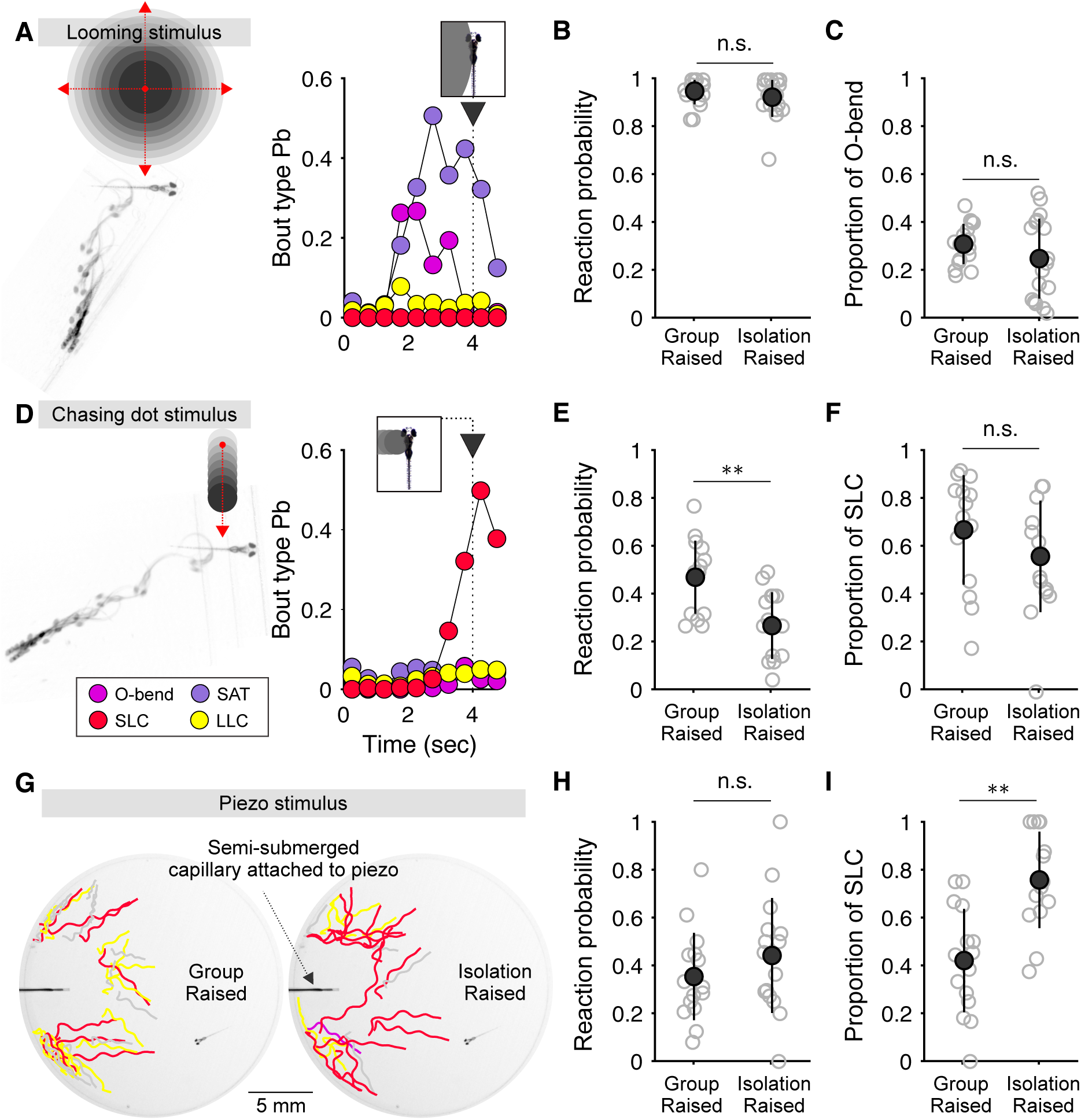
Avoidance bout type responses upon controlled closed-loop stimuli. (A) A radially expanding dark spot (1 cm/s) was projected 4 cm away from the larva. The outer edge of the stimulus reaches the larva’s center of mass after 4 sec (shown as inset). As the dark spot expands, the likelihood of performing O-bend and SAT bout types increases, but not SLC and LLC. (B) The probability of performing an O-bend or SAT during the visual stimulation period per stimulus presentation was calculated for each larva tested for group and isolation raised larvae (*N*_GR_ = 16, *N*_ISO_ = 17, *p* = 0.3209). (C) The proportion of O-bend responses calculated as the number of O-bends performed during looming stimulus presentation divided by the total number of stimulus-induced O-bends and SATs (*N*_GR_ = 16, *N*_ISO_ = 17, *p* = 0.3678). (D) A non-expansive dark spot (0.5 to 2.5 mm in diameter) was projected at 2 cm away from the test larva and approached at a speed of 0.5 cm/s. Hence, the outer edge of the stimulus reaches the center of mass of the larva after 4 sec (shown as inset). Such stimulus increases the probability of performing SLCs. (E) Reaction probability is defined as the number of SLC or LLC performed per total number of chasing dot stimulus presentations per larvae or either raising condition (*N*_GR_ = 14, *N*_ISO_ = 15, *p* = 0.0022). (F) The proportion of SLCs was calculated as the number of stimulus induced SLCs divided by the sum over stimulus-induced SLCs and LLCs (*N*_GR_ = 14, *N*_ISO_ = 15, *p* = 0.2384). (A,D) Shown are the overlaid image sequences of an O-bend (A) and an SLC (D) performed by a group raised larvae, acquired at 90 frames per second. The bout type probability was calculated as the number of bouts of a given bout category performed within a 500ms time window divided by the total number of bouts performed. (G) Local water vibrations per elicited by submerging the tip of a rod into the swimming area. The rod was attached to a piezo bender which would trigger a pulse train of five 2ms pulses with an input voltage of 40V at a minimum inter-stimulus interval of 2 min. Stimuli were triggered when the larva was facing away from the source at a distance of 5mm from the tip of the rod. Shown are the trajectories of the first bout after piezo stimulus presentation for five randomly chosen group raised (left) and isolation raised larvae (right). (H) Reaction probability is defined as the number of SLC or LLC performed per total number of chasing dot stimulus presentations per larvae or either raising condition (*N*_GR_ = 15, *N*_ISO_ = 15, *p* = 0.7714). (I) The proportion of SLCs was calculated as the number of stimulus induced SLCs divided by the sum over stimulus-induced SLCs and LLCs (*N*_GR_ = 15, *N*_ISO_ = 15, *p* = 0.0017). (B,C,E,F,H,I) Light gray open circles show every replicate per condition, dark gray filled circles and error-bars signal mean ± standard deviation. Asterisks specify the results of unsigned Wilcoxon tests between raising conditions, alpha level 0.05. SAT: Shadow avoidance turn; LLC: Long latency C-start; SLC: Short latency C-start.

To test the contribution of the LL to social avoidance reactions, we designed a stimulus to mimic the water disturbances caused by a swimming larva. While sudden acoustic stimuli are effective at triggering C-start responses, they lack a directional component and are quite unlike a social stimulus. De Marco et al. [2013] showed that local water vibrations can act as a stressor for larval zebrafish and enhance their level of cortisol in an intensity-dependent manner. To cause these vibrations, a rigid capillary was attached to a piezo bender and semi-submerged into the water of the swimming arena. We replicated a similar setup and programmed the triggering of the piezo displacements in closed-loop with the swimming pattern of the larva. In order to mimic the conditions during social avoidance reactions, piezo displacements were triggered when the larva was facing away from the stimulus source at a distance of 5mm. To avoid triggering the stimulus as the larva is performing a bout, we added a 400ms delay between the crossing of the distance threshold and the onset of the piezo. This stimulus protocol was effective as triggering both types of C-starts (Fig. 4 G; Fig. S4 C) in GR and ISO larvae. While both raising conditions were comparable in their overall response probability (Fig. 4 H, *p* = 0.25), ISO larvae showed a higher probability of using SLCs (Fig. 4 I, *p* = 0.0016, see Table S10), matching our findings during social interactions. Notably, the latency profile of LLC and SLC upon piezo stimuli remains consistent with the naming of the two C-starts, although they are not as clearly separated as with acoustic startle stimuli (Fig. S4 E-F). This resembles our findings of LLC and SLC latency during social interactions.

To validate that the piezo-induced water motions are indeed a LL stimulus we measured the responses of GR larvae after neomycin treatment to ablate the LL hair cells. The overall response probability (*p <* 0.0001), as well as the proportion of SLC responses (*p* = 0.0002; Fig. S4 G-I, see Table S10) were significantly reduced although not entirely abolished with the ablation. These results show that LL sensing is the main, albeit not sole source of processing for water motion stimuli. Overall, we here showed that LL dependent stimuli, composed to local water vibrations, recapitulate the phenotype of social isolation raising during social avoidance reactions. Hence, indicating that LL sensing is sufficient to drive the differences between GR and ISO.

## Discussion

Social experiences during early life greatly impact the development of social and non-social behaviors [Harlow et al., 1965]. To date, it is not fully understood what the underlying mechanisms are by which social experiences shape appropriate behavior and development. Here we set up to test if social isolation effects can be studied in larval zebrafish whose social experiences can be precisely controlled and whose social behavior can be studied as discrete, quantifiable events. We found that isolation raising enhances LL mediated avoidance reactions to another fish, along with general effects on locomotion.

### The effects of social isolation on social interactions can already be observed in one-week-old larvae

Previous studies have reported that social experience is not necessary for the development of social attraction in zebrafish [Breder and Halpern, 1946, Larsch and Baier, 2018]. However, social isolation led to reduced group cohesion and a larger average distance between groups of freely swimming adult zebrafish [Shams et al., 2018]. In line with this, we found that the avoidance distance is larger in larval zebrafish after isolation raising. To the best of our knowledge, we here showed for the first time that the effects of social isolation can already be observed during social interactions in one-week-old zebrafish larvae when social attraction is yet to be developed. Together these results suggest that social attraction develops without the need for social experience, while fine-tuning locomotion for regular collective motion requires the presence of conspecifics during development.

### Isolation affects general locomotion

We also observed that ISO larvae have general, albeit small, deficits in locomotion. They produce longer and fewer bouts than GR larvae, irrespective of the social context during testing. Hypoactivity in isolation raised zebrafish larvae has previously been reported upon exposure to a sudden dark period during the larva’s day time [Zellner et al., 2011, Steenbergen et al., 2011]. While our findings are in line with these results, there is a noteworthy difference. We here tested spontaneous locomotion with continuous background illumination, while in the referred publications larvae were dark-adapted and then tested in alternating light patterns. Exposing a dark-adapted larva to sudden changes in illumination leads to increased levels of the stress hormone, cortisol [De Marco et al., 2013]. Therefore protocols using light dark alternation may test specifically for locomotion under acute stress. A previous study has reported increased locomotor activity in juvenile, but not larval zebrafish after isolation raising [Shams et al., 2018]. We have observed a similar effect in a pilot experiment, when raising ISO fish to three weeks of age (data not shown). However, we also noted that ISO fish were growing faster than GR fish once they were fed, thus the change in direction of the isolation effects on locomotion with age, may result from an interaction with feeding. Nonetheless, general effects of social isolation on locomotor activity have been observed in several species and can vary in sign and magnitude (e.g. Wilkinson et al. [1994] and Archer [1969]). This further strengthens the point that when studying the impact of social isolation on social behavior, it is essential to thoroughly distinguish general locomotion and social-specific effects.

We here designed our experiments to parse general effects on spontaneous locomotion apart from social effects of isolation raising, by comparing locomotion parameters across different social testing contexts and isolation time windows, as well as manipulating locomotion pharmacologically. Therefore, two lines of evidence in our data support the hypothesis that general and social locomotor effects are distinct. First, the increased bout duration and social avoidance phenotype have dissimilar critical developmental windows. Isolating larvae at the third day of development abolishes the general increase of bout duration, but does not detract the behavioral phenotype during social avoidance. Secondly, treatment with the generic dopamine agonist, apomorphin, recapitulates the general locomotion of ISO fish in GR fish without affecting the social avoidance measures.

### Isolated fish show enhanced use of SLCs in social context

An advantage of using one-week-old zebrafish is that the repertoire of their swims has been described and can be clustered into movement types through automatic, unsupervised methods [Marques et al., 2018, Johnson et al., 2020, Mearns et al., 2020], enabling exact phenotyping of behavioral events. Using this technology, we found that larvae avoid each other by executing four types of large displacement bouts: burst swims [Severi et al., 2014], long capture swims [Marques et al., 2018], O-bends [Burgess and Granato, 2007b], and C-starts (LLCs and SLCs) [Burgess and Granato, 2007a]. Strikingly, although the use of C-starts as a whole is unaltered, ISO fish use SLCs twice as frequently as LLCs when compared to GR fish. SLCs are a high acceleration version of the C-start and lead to a larger displacement. Therefore, the imbalance in usage of the two C-start types could be a major contributor to the enhanced avoidance area seen in ISO larvae. Given the larger displacement, SLCs increase the interfish distance faster than LLCs. In line with this argument, after ablation of the lateral line, a manipulation that balances the use of C-starts, the avoidance area difference between raising conditions disappears.

### Larval social avoidance is multimodal and isolation impacts lateral line mediated C-starts

The social stimuli during collective swimming are multimodal and fish rely mostly on vision and the LL. Only upon removal of both of these, fish fail to swim in a coordinated fashion [Pitcher et al., 1976, Partridge, 1982]. It has been proposed that the contribution of these two sensory systems to schooling is distinct; vision is thought to be primarily important for maintaining position and angle between neighboring fish, while the LL contributes to monitoring swim speed and direction of swimming neighbors [Partridge and Pitcher, 1980]. Presumably due to do the technical difficulty of experimentally mimicking water wakes of a swimming fish, to date only visual stimuli have been shown to be sufficient as a sole motion cue to induce following behavior in adult zebrafish [Lemasson et al., 2018], as well as juvenile zebrafish [Larsch and Baier, 2018]. The social stimuli that trigger social avoidance in larval zebrafish are also multimodal and previous work has shown that vision plays a critical role. First, the avoidance area is decreased when fish are interacting in the dark [Marques et al., 2018], suggesting that vision helps larvae avoid each other at larger distances. Secondly, through visual inputs larvae and juvenile fish synchronize their movements with other fish [Dreosti et al., 2015, Marques et al., 2018, Stednitz and Washbourne, 2020].

To parse apart the contribution of vision and LL sensing, we tested larvae with manipulations that selectively blocked each modality. While testing social avoidance in darkness has a strong effect on the avoidance distance, it has no impact on the C-start choice, suggesting that vision defines the avoidance distance, but not the type of swim bout used. The avoidance distance in darkness is similarly low for GR and ISO fish, pointing towards mixed effects of isolation raising and the availability of visual cues. The LL, on the contrary, seems to contribute specifically to the effect of isolation on the balance of the two C-start types. In line with this, we did not observe enhanced reactions to visual stimuli. To probe sufficiency of mechanosensory stimuli, we designed a social-like LL stimulus inspired by previous work that used a piezo bender to produce local water vibrations [De Marco et al., 2013, Groneberg et al., 2015]. The stimulus parameters here were chosen to mimic the conditions we observed during social escape reactions in freely interacting fish. To ensure that all larvae receive comparable sensory input, we triggered the stimuli in closed-loop with the behavior of the testing larva. Thereby we found that ISO fish also showed an enhanced proportion of SLC usage. This assay was performed in darkness, but we additionally confirmed through neomycin treatment that it requires the LL. Hence, LL stimuli are sufficient to trigger enhanced social avoidance reactions in ISO larvae.

Notably, it has previously been reported that during the acoustic startle assay, larvae raised at a lower density are more likely to perform SLCs reactions [Burgess and Granato, 2008]. Generally, this report is in accordance with the results shown here. However, we believe that the social isolation effects during social avoidance represent a different scenario. Through the ablation experiments we learned that the LL sensing is necessary for the increased SLC proportion in ISO larvae. Acoustic startles mainly depend on the inner ear and not the LL [Lacoste et al., 2015] and are strong, non-localized stimuli, unlike the water disturbances caused by another larva swimming in proximity. Accordingly, we also observed differences in the latency profile of the C-starts between acoustic and piezo stimuli. This suggests that although the bout types are kinematically the same, the responses to minute water vibrations, artificial or socially-borne, are differently triggered from the highly stereotyped acoustic startle reactions.

### Neural control of startle reactions and the social isolation phenotype

The neural circuits underlying escape reactions in larval zebrafish have been studied for over two decades [Eaton et al., 1991, Liu and Fetcho, 1999, O’Malley et al., 1996, Kohashi and Oda, 2008, Burgess and Granato, 2007a]. Most of the studies have used acoustic or tactile stimuli upon which the neural populations controlling SLCs and LLCs appear to be distinct. The prominent Mauthner escape neurons (M-cells), a pair of large cells in the dorsal hindbrain segment (rhombomere) 4 [Kimmel et al., 1981], have been studied in great detail and were shown to be necessary for SLCs but not LLCs [Burgess and Granato, 2007a, Takahashi et al., 2017]. More recently, a study identified a cluster of prepontine neurons, in hindbrain rhombomere 1, as necessary and sufficient to drive LLCs [Marquart et al., 2019]. In line with their delayed response outputs, these neurons neither project directly to the spinal cord, nor receive direct auditory input. Moreover, prepontine neurons are only active when the larva performs an ipsilateral LLC, but not when it performs an SLC, suggesting that the circuits driving the two C-start types are active in a mutually exclusive manner [Marquart et al., 2019].

M-cell excitability is regulated through a network of excitatory and inhibitory feedback and feed-forward interneurons [Takahashi et al., 2002, Satou et al., 2009, Koyama et al., 2011, Lacoste et al., 2015, Shimazaki et al., 2019]. Behavioral switching from SLC to LLC during conditioning or habituation is associated with reduced excitability of the M-cell [Takahashi et al., 2017, Marsden and Granato, 2015]. Furthermore, forward genetic screens have shown that mutants of altered SLC-LLC reaction balance are associated with altered M-cell excitability [Marsden et al., 2018, Jain et al., 2018].

Therefore, one explanation of why ISO larvae perform more SLCs during social interactions, is an increased M-cell excitability. However, in this case, one would expect to also see enhanced responses to other stimuli that trigger M-cell dependent escapes, such as the approaching dark spot [Dunn et al., 2016] and acoustic startles [Burgess and Granato, 2007a]. An alternative mechanism would be that the LL in ISO larvae is more sensitive to the water motions caused by a conspecific swimming in proximity, as these represent a novel stimulus. Although it has been shown that the connections between afferent neurons and hair-cell bundles of the LL neuromast develop independently of experience in larval zebrafish [Nagiel et al., 2009], this sensory organ shows several filtering properties whose development might depend on experience. For instance, efferent copies of forward motion suppress synaptic transmission in lateral line hair cells [Pichler and Lagnado, 2020], and canal neuromasts are capable of sensing water vibrations even against a background of unidirectional water flow [Engelmann et al., 2002]. Such fine-tuning of LL sensing under specialized conditions could be shaped through the sensory experiences of the presence of conspecifics.

In support of this latter hypothesis, it has previously shown that a hypoactivity response to subtle, minute water vibrations was more sensitive in larvae that were raised in isolation [Groneberg et al., 2015]. Notably, in this behavioral assay the larvae did not perform an escape reaction to the stimulus presentation and therefore an enhanced response indicates higher sensory sensitivity rather than reduced startle thresholds.

To fully probe these two possible mechanisms of enhanced social avoidance reactions in ISO larvae, future experiments will need to measure neural activity in the LL ganglion cells, as well as the M-cells upon controlled stimuli of graded intensity, along with social stimuli in larvae where the LL is free and intact.

## Conclusion

Understanding how the brain implements adequate, fine-tuned social behaviors according to social experiences, requires precise experimental control of such experiences along with detailed behavioral analysis. Another critical choice is a model that will allow studying ethologically relevant social interactions, along with the capacity to measure and manipulate activity in neural circuits. Here we showed that a relatively simple social behavior, the avoidance reactions to a minute, local social-like stimuli, is shaped by experience and therefore more complex than previously thought. Moreover, we show that the effects of early life social isolation can be distinguished between general locomotor effects and social behavior, at a young age in a model which allows whole-brain neural activity measure and manipulation.

## Author contributions

AHG, MBO and GGdP conceived the project. AHG and JCM designed experiments. JCM and MBO implemented the recording apparatus and AHG performed all experiments. ALM programmed the piezo stimulus. AHG and JCM analyzed data and all authors interpreted the results. AHG and JCM wrote the manuscript with contributions from all authors. MBO and GGdP supervised the project.

## Acknowledgements

We thank Alexandre Laborde for assistance in setting up the experiments, and Mattia Bergomi and Sabine Renninger for discussions on the project. We also thank the Champalimaud Fish Facility team for excellent fish care, and Paulo Carriço and the Champalimaud Hardware platform for logistic support.

## METHODS

### Experimental model and subject details

Zebrafish breeding and maintenance was performed under standard conditions [Westerfield, 2007, Martins et al., 2016]. The sex of the larvae is not defined at this early stage of development. Fertilized wild type embryos of the Tubingen (TU) strain were collected in the morning and placed in petri dishes containing E3 medium [Westerfield, 2007]. Embryos were kept in 10 cm petri dishes in groups of 20 for the first three days of development. Then at 3 dpf, larvae were randomly assigned to a raising condition (isolation raised or group raised) and placed either alone or in groups of 8 into 35 mm petri dishes containing fresh E3 medium. All dishes were kept at 28°*C* and a 14/10 h light-dark cycle, starting at 8 am. Larvae were tested at 6 dpf. All behavioral experiments were performed in a randomized order during the larva’s day period, between 9 am and 8 pm. Animal handling and experimental procedures were approved by the Champalimaud Center for the Unknown Bioethics Committee and the Portuguese Veterinary General Board (Direção Geral Veterinária), in accordance with the European Union Directive 86/609/EU.

### Behavioral recording apparatus

The behavioral setup has previously been described by [Marques et al., 2018]. Inside of a light-proof enclosure, a high-speed infrared sensitive camera (MC1362, Mikroton) with a Schneider apo-Xenoplan 2.0/35 lens and a 790 nm long pass filter was mounted above a stage holder for the fish recording chamber. Larvae were illuminated from below by a 10 x 10 cm infrared LED-based diffusive backlight (850 nm). Tracking of the larva’s position and equally spaced tail segments of 0.3 mm was performed online during video acquisition at 700 frames per second using a custom-made algorithm in C#, as previously described by [Severi et al., 2014, Burgess and Granato, 2007a]. The tail data was interpolated to a standardized set of points in the tail spaced at 310 *µ*m increments, starting at a point 1345 *µ*m behind the midpoint between the two eyes.

### Swim bout detection

We applied the bout detection algorithm described in [Marques et al., 2018]. To detect bouts, a measure of change in tail curvature was used to capture tail movements reliably. To compute this value, we applied spatiotemporal smoothing to the raw tail angles using a boxcar filter (3 segments, 10 frames) and calculated the frame-to-frame change of this smoothed measure at each point along the tail. For each segment, we then calculated the cumulative sum (from rostral segment to current segment), took the absolute value, and then summed the results over all segments to obtain a scalar measure of tail curvature. This helps to ignore local tail fluctuations and highlights extended regions of curvature. To exclude effects of slow drifts in tail position, we subtracted the minimum of the previously obtained curvature measure over a 571 ms window. To smooth out the half-beats that compose individual bouts, we applied a maximum filter with a 30 ms window. For each fish the maximum of this measure was calculated and divided by 20, and this value was used as a threshold to detect individual bouts. Short fluctuations of the tail curvature measure that did not correspond to tail movements were excluded by ensuring that bouts are at least 43 ms long. To ensure that individual bouts were not broken into parts, the interbouts must also have a minimum duration of 43 ms. The exact start and end of bouts was calculated by the start of the first half-beat and the end of the last half-beat (see half-beat detection).

### Half-beat detection

Half-beats were detected as previously described [Marques et al., 2018]. A dynamic threshold was computed for each bout by using a 43 ms boxcar filter to smooth the tail curvature values. The beginning and end of each half beat, at each segment, were found by locating the points where the tail crosses this threshold. The half-beat peak at each segment was defined as the maximum absolute value of tail angle. Some large amplitude bouts had a first half-beat that was a small passive deflection in the opposite direction of the second larger half-beat. These situations are detected by finding small first half-beats (10% higher than the baseline value) that were followed by larger movements in the opposite direction. To maintain the consistency of the half-beat numbering, for these bouts, the first small half-beat was excluded and the second larger half-beat was considered the first.

### Bout type categorization

For each swim bout, we computed 73 kinematic parameters based on the bout and half-beat detection, as described previously in depth [Marques et al., 2018]. After, we embedded all the data into a previously computed principal component analysis (PCA) space and included for further analysis the first 20 Principal Components (PCs). This space was obtained by performing PCA over the 73 kinematic parameters on a dataset with 3 million bouts, that fish executed while performing 9 behaviors over 255 stimuli, that included social avoidance in the light and dark, acoustic startles, and responses to looming stimuli [Marques et al., 2018]. Therefore, we expect that the new data gathered in this report, that is mainly constituted of avoidance bouts, to be well represented in this PCA space. To categorize the bouts into types, we used a dataset composed of equal number of bout types (Bout Map) [Marques et al., 2018], that were categorized using density valley clustering [Marques and Orger, 2019], and used k nearest neighbors (k = 50) to assigned every new bout to one of the 13 possible bout categories.

### Behavioral testing of spontaneous locomotion

Larvae were carefully placed in a circular swimming arena with a depth of 3mm inside the recording setup. After 5 min of adjustment time the recording was started and lasted for 1 hour. Pairs of larvae of the same raising condition were tested in circular arenas with a 20 mm diameter. One recording chamber consisted of four arenas, hence eight larvae could be recorded at once. All arenas had a transparent bottom plate and black acrylic walls to avoid that larvae could see the pairs in the neighboring arenas. For recordings in light, a homogeneous background illumination of 1000 lux was projected onto an opal glass diffuser 5 mm below the larvae by a DLP projector (BenQ). For recordings in darkness, the projector remained turned off, hence the illumination inside the light-proof enclosure was 0 lux.

### Avoidance area measure

The avoidance area measure was adjusted from [Marques et al., 2018]. First, we computed the density distribution of the positions of the other larvae relative to a focal larva for each frame of the experiment. Then we created a reference density distribution by randomizing the order of the recording frames. Both, the data and the reference density distributions were then smoothed by a Gaussian filter with a kernel sigma of 0.36mm. To avoid arbitrarily high ratio values due to division by a value near zero, points at the edges where the randomized control data had a value less than 1 were excluded. The raw density data was divided by the control to generate a ratio image. The low density region in the center of the image was quantified by counting the pixels in a bounded region where the density of the real data was below 60% of the randomized control density. The contours of these detected areas are shown in the right panel of Fig. 1B.

When testing pairs of larvae instead of groups of seven, the data is more sparse and the thresholding of the density distribution can lead to false high values in area. We therefore adjusted the avoidance measure to a one-dimensional version by considering the distance between the two fish rather than a two-dimensional map of positions. Similarly to the 2D avoidance area measure, we created a random distribution by shuffling the frame order so that the trajectories of both fish were no longer matched in time. We then made a histogram of the values of the real and the randomized data with a bin size of 0.5*mm*. The ratio between the randomized and the real data histogram was negative for close distances, indicating that the fish avoid each other. To quantify this avoidance we used a trapezoidal numerical integration of the distances smaller than the zero crossing.

### Comparison of SLC and LLC

The proportion of SLCs is calculated by dividing the total number of SLCs performed by the sum over all C-starts (SLC and LLC) performed when the two fish were swimming within 10*mm* of one another. Position of the non-focal fish upon C-start onset of the focal fish is considered for when the distance between the two fish was smaller than 10*mm* and the nonfocal fish performed a bout within 1000*ms* prior to C-start onset.

### Cluster comparison between C-starts of group and isolation raised fish

The LLC and SLC bouts were embedded in the same PCA space described in the “Bout categorization” section and the first four PCs were taken into account. If raising conditions modulate kinematically the bout types, we expect them to occupy a distinct place in this space. For visual comparison we plotted 200 randomly chosen LLCs and SLCs per raising conditions into the first four components of the PCA space.

### Pharmacology

The general dopamine agonist, apomorphine (R-(-)-Apomorphine hydrochloride hemihydrate, Sigma-Aldrich, A4393), was prepared to a final concentration of 20 *µ*M in buffered E3 medium. Larvae were incubated for 10min in the drug, and tested after three washes and a recovery period of 10min. This protocol is in line with what has been published in adult zebrafish [Stednitz et al., 2018].

For lateral line ablation, *CuSO*_4_ (Copper(II) Sulfate pentahydrat, Acros Organics) was dissolved as concentration of 1 mM in E3 medium. Larvae were incubated for 1h without a change in social context; isolation raised larvae were incubated in isolation and group raised larvae in groups. After incubation larvae were washed three times in fresh E3 medium and given a 1h recovery period prior to behavioral testing. Neomycin trisulfate (Sigma-Adrich) was dissolved in tris-buffered E3 medium (E3-tris, 1 mM tris) to a final concentration of 200 *µ*M. Larvae were incubated as described above for 1h. Thereafter larvae were washed three times in E3-tris and given a 2h recovery period before behavioral testing.

Both of these incubation protocols have been shown to ablate hair cells with a regeneration time larger than our testing time window [Buck et al., 2012].

Handled controlled for *CuSO*_4_ and neomycin treatments were exposed to the same treatment with incubation in E3 medium or E3-tris, respectively.

### Closed-loop presentation of visual stimuli

Single larvae of either raising condition were swimming freely in a circular arena (5 cm in diameter) with a homogeneous background illumination projected onto an opal glass diffuser 5 mm below the larvae by a DLP projector (BenQ). Both types of visual stimuli were presented with an inter-stimulus interval of 2 min. The walls of the arena were transparent, so that visual stimuli could be displayed even when the larva was swimming in the arena borders. For testing looming stimuli an expanding dark spot (1 cm/s) was projected 4 cm away from the larva, either placed to the front, the back, left or right of the larva [Marques et al., 2018]. In the approaching dark spot assay a spot of constant size was projected 2 cm away from the larva and approached it with a constant speed of 0.5 cm/s. The spot radius varied between trials from 0.5 to 2.5 mm and was approaching from the front, the back, left or right. All stimuli were tested twice per animal and presented in a randomized order. Each randomized order was presented to one group raised and one isolation raised larva to avoid any unbalanced order effects.

### Closed-loop presentation of mechanosensory stimuli

To produce minute local water displacements we used a piezoelectric bender actuator (Thorlabs; voltage range: 150V; max. displacement: 450 *µ*m). A 2 cm long capillary rod was attached to the body of the piezo bender using silicone. The rod was made of the tip of a loading-pipette tip (outer diameter: 0.3 mm). To give more rigidity to the rod a headless and stainless steel insect pin was pushed inside. The larger end of the capillary was melted into a flat circle via a heat-shrink gun to allow for better attachment to the piezo bender. The capillary rod attached to the piezo bender was mounted vertically at a 45° angle, so that 2 mm of its tip would be submerged into the water of the swimming arena. We left a single larva of either raising condition swim inside this arena for one hour in complete darkness. Movements of the capillary tip were triggered in closed-loop depending on the position and movement of the larva with a minimum interval between capillary stimuli of 2*min*. We used the following criteria to trigger piezo deflections: 1) the larva had to be less than 4*mm* away from the tip of the capillary rod; 2) thereafter the larva had to be more than 5*mm* away from the tip; 3) after a 400*ms* delay the piezo was triggered. Unless otherwise stated the structure of the piezo pulse was composed of five 2*ms* pulses with a 2*ms* inter-pulse interval. The voltage applied stayed fixed at 40*V* for each pulse train. Each fish was only exposed to one set of fixed stimulus parameters. Different sets of stimulus parameters were tested across different fish. Testing times and fish clutches were interleaved to avoid any uncontrolled bias.

### Acoustic startle assay

A single larva was swimming for 1 hour and 40 min in a custom-build chamber. with a circular swimming arena of 5 cm diameter. Outside of the swimming arena was a black acrylic sheet onto which four speaker membranes (Visaton) were glued with equal spacing. A single 100 ms pure tone of 600 Hz was played by gating the sound output of the recording PC through a digital switch (Arduino One). Acoustic stimuli were presented with a minimum ISI of 6min. The experiment was interleaved with other visual stimuli, which were not analyzed for this manuscript. In total 16 acoustic startle stimuli were presented per larva.

### Quantification and statistical analysis

All data are either presented as pooled events per raising condition (when plotted as histograms), or as single measure per replica and shown as mean with standard deviation, unless otherwise stated in the figure legend. Only datasets that were collected in the same days of experimentation are considered for statistical testing. Datasets composed of pairs or groups of larvae per recording were first averaged between individuals and then this average was considered as the unit of comparison across replicates, so as to avoid pseudo-replication. We applied two-tailed unsigned Wilcoxon tests (Matlab: ranksum) for comparing between raising conditions or treatments and two-tailed signed Wilcoxon tests (Matlab: signrank) when comparing repeated measures coming from the same larva or pair of larvae, i.e. comparison between bout types. When performing series of pairwise comparisons, p-values were corrected for the number of comparisons using Holm’s sequential Bonferroni procedure (www.mathworks.com/matlabcentral/fileexchange/28303-bonferroni-holm-correction-for-multiple-comparisons). For comparing the distribution of angles of the position of the non-focal larvae upon C-start onset, we used the CircStat toolbox [Berens, 2009]. As an addition to regular hypothesis tests, for all comparisons between raising conditions or treatments, we quantified the effect sizes of the observed differences using the Measures of Effect Size (MES) Toolbox [Hentschke and Stüttgen, 2011]. In the statistical summary tables in the supplementary material, as well as in Fig. 3J we used the mean differences and its confidence intervals as calculated with 1000 bootstraps. All analysis and statistical tests were performed in Matlab 2018a.

## Supplementary Material

**Figure S1:**
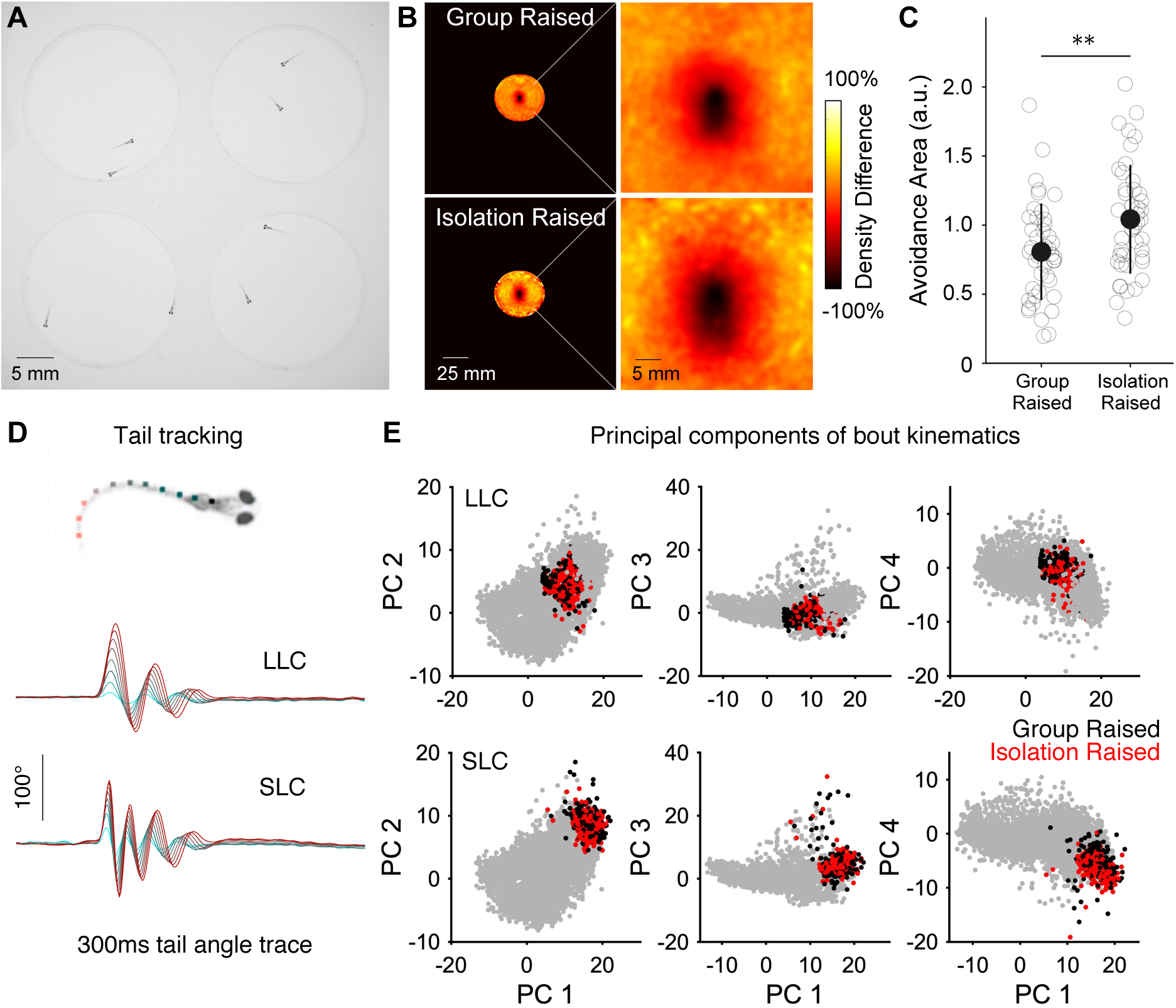
Social avoidance area measures in pairs of larvae. (A) Pairs of larvae raised either in groups or in isolation were video recorded for 1 h while swimming in a circular arena with a 20*mm* diameter. (B) Density distributions, relative to a focal individual, of group raised (top) and isolation raised larvae (bottom). The focal fish is located in the center and facing upwards. Raw densities are divided by a reference distribution created by choosing fish locations from random frames. Right panels are zoomed views of the black squares in the left panels. (C) For pairs of larvae the avoidance distance was calculated as the negative area under the curve of histograms comparing the distance between the two fish to a control distribution of randomly shuffled recording frames. Light gray open circles show every replicate per condition, dark gray filled circles and error-bars signal mean ± SD. Asterisks specify the results of a two-tailed unsigned Wilcoxon test; p ¡ 0.001; N=43). (D) Tracking of equally spaced tail segments at 700 Hz enables measuring the tail oscillations of each swim bout. Shown here are example traces of tail angles across a 300 ms sequence containing a long latency C-start (LLC, top) and a short latency C-start (SLC, bottom). The measures of all tail segments are overlaid and color-coded according to the position along the body of the fish show on the top. (E) Mapping of 200 randomly selected LLCs (top) and SLCs (bottom) for group (black) and isolation-raised larvae (red), in the space of the first four principal components that were used for the bout classification based on 73 kinematic parameters. Gray scatter points display all bouts that are not C-starts. Group and isolation data points were randomized so that the colors are not overlaid.

**Figure S2.**
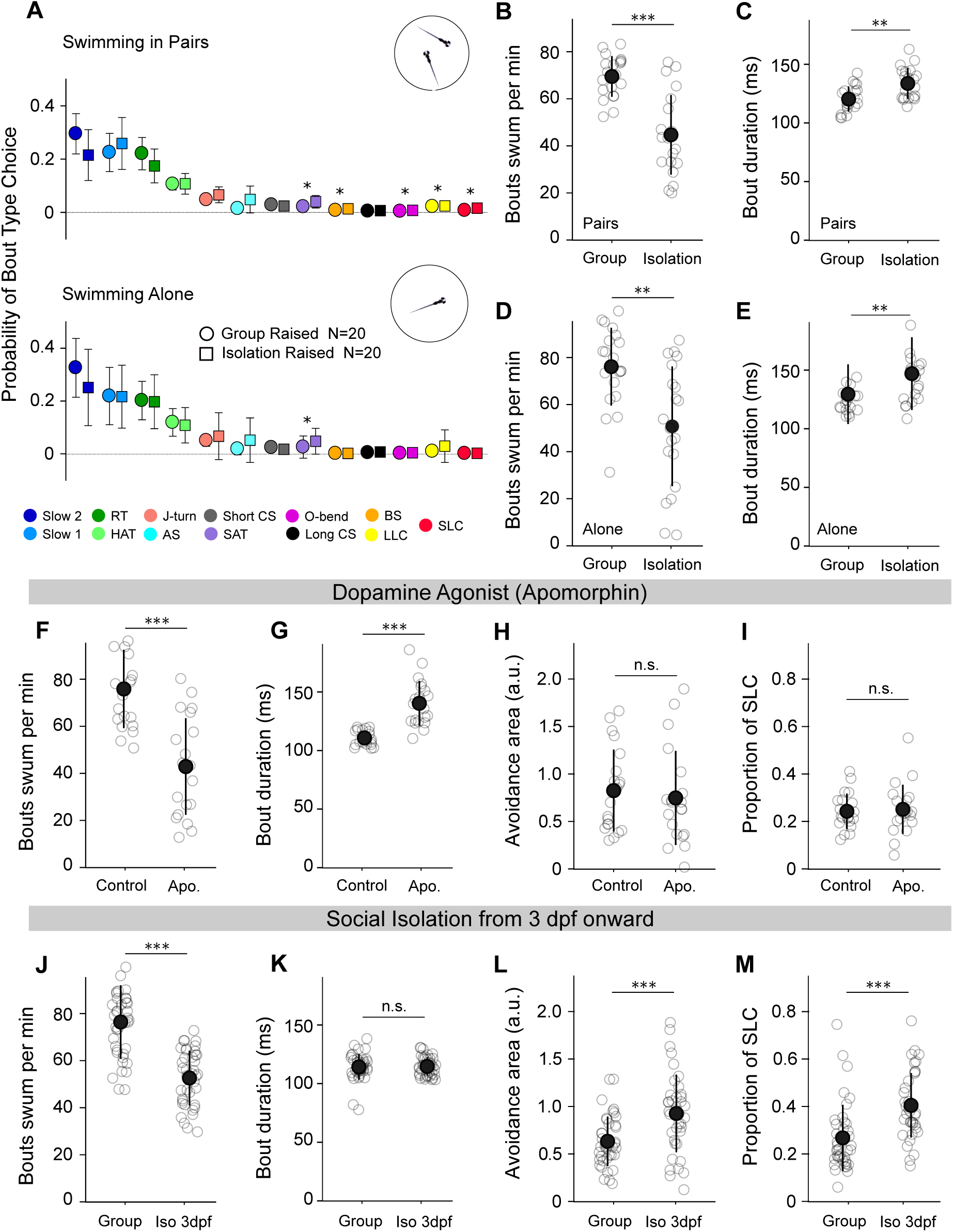
General locomotion effects of isolation raising. (A) Bout type probability during 30min of spontaneous locomotion of larvae tested in pairs of the same raising condition (top) or swimming alone in the arena (bottom). Bout types are color coded, markers and errorbars mean ± SD, for group raised (circles) and isolation raised larvae (squares). Asterisks signal significant results of bonferroni-holm corrected p-values from two-tailed unsigned Wilcoxon tests for each bout type (alpha level 0.05). (B,D) Number of bouts swum per min over the 30 min recording period (N=20). (C,E) Average duration of each bout performed irrespective of the bout category (N=20). (F-I) Group raised larvae were treated for 10min with the generic dopamine agonist apomorphin (Apo.) (N=20). (J-M) Instead of a full-term isolation protocol, larvae were raised in groups until 3dpf and thereafter isolated (Iso 3dpf) or handled similarly and placed in a new dish remaining in the social context (Group) (*N*_GR_ = 40, *N*_ISO_ = 39). Larvae were tested in pairs of the same treatment group. Shown are the measures of number of bouts swum per min over the 30 min recording period (F,J), the average bout duration (G,K), the avoidance area as calculated by comparing the distribution of inter-fish distances with a random distribution (see methods; H,L) and the proportion of SLC out of all C-starts (LLC and SLC) performed as the larvae swam within 10mm of one another (I,M). Light gray open circles show every replicate per condition, dark gray filled circles and error-bars signal mean ± SD. Asterisk specify the results of statistical comparisons between raising conditions. AS: Approach Swim; Long CS: Long capture swim; BS: Burst swim; HAT: High angle turn; RT: Routine turn; SAT: Shadow avoidance turn; LLC: Long latency C-start; SLC: Short latency C-start.

**Figure S3:**
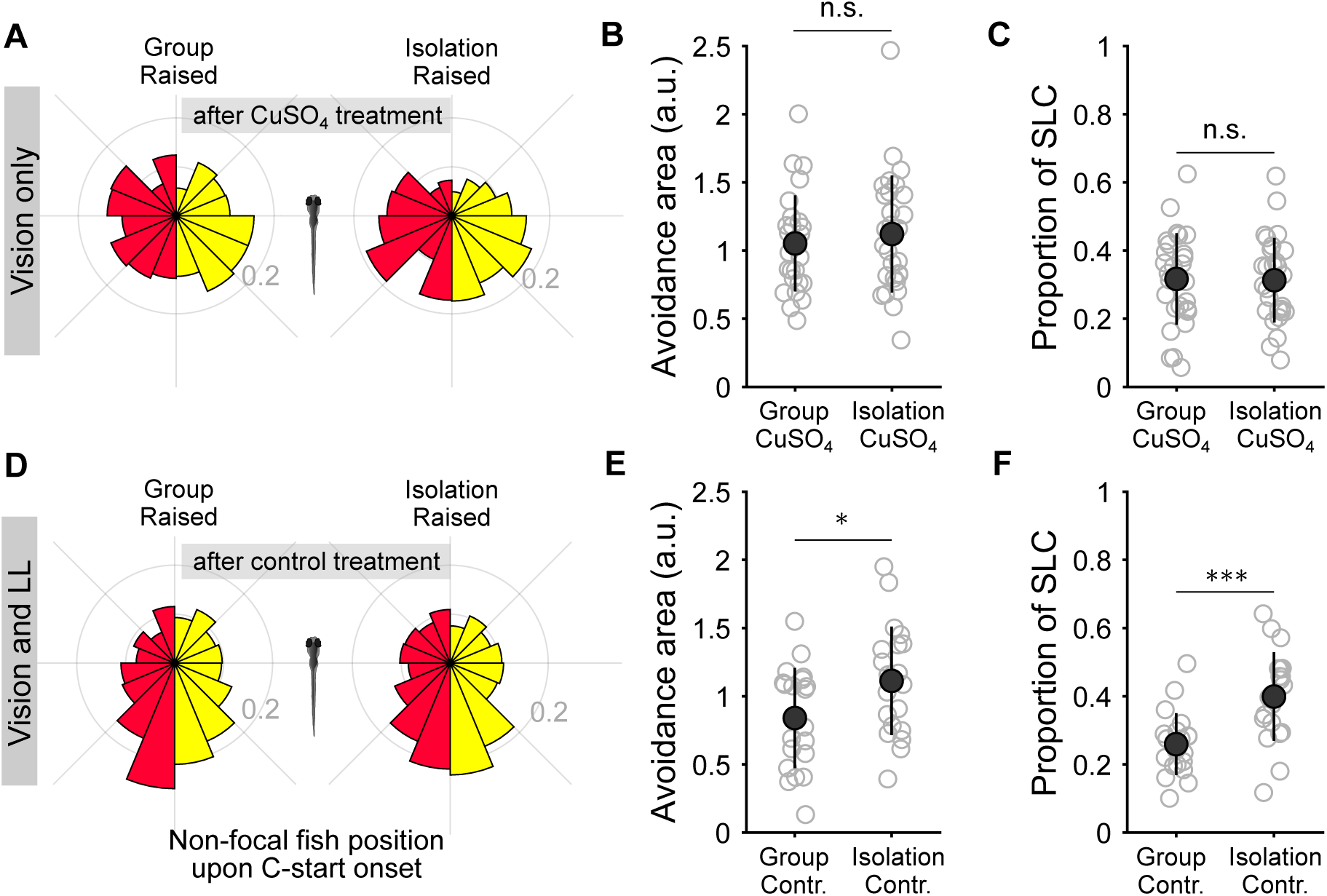
Effects of *CuSO*_4_ and control treatment of social avoidance reactions. (A-C) Larvae were incubated for 1h in *CuSO*_4_ and given a 1 hour recovery period prior to testing with homogeneous background illumination; *N*_GR_ = 27; *N*_ISO_ = 28. (D-F) Control treatment consisted of incubating larvae for 1h in regular E3 medium, performing the washing steps and then allowing for a 1 hour recovery period prior to testing with homogeneous background illumination; *N*_GR_ = 20; *N*_ISO_ = 20. (A,D) Circular histograms of the position of the non-focal larva upon the onset of the C-start. Data is pooled over fish per raising condition and treatment. Red and yellow bars show data for SLC and LLC, respectively. (B,E) Social avoidance area as calculated from the distribution of distances between the two fish throughout the recording period. (C,F) The proportion of SLC bouts over the sum of SLC and LLC. Light gray open circles show every replicate per condition, dark gray filled circles and error-bars signal mean ± SD. Asterisk specify the results of two-tailed unsigned Wilcoxon tests between raising conditions (alpha level 0.05).

**Figure S4.**
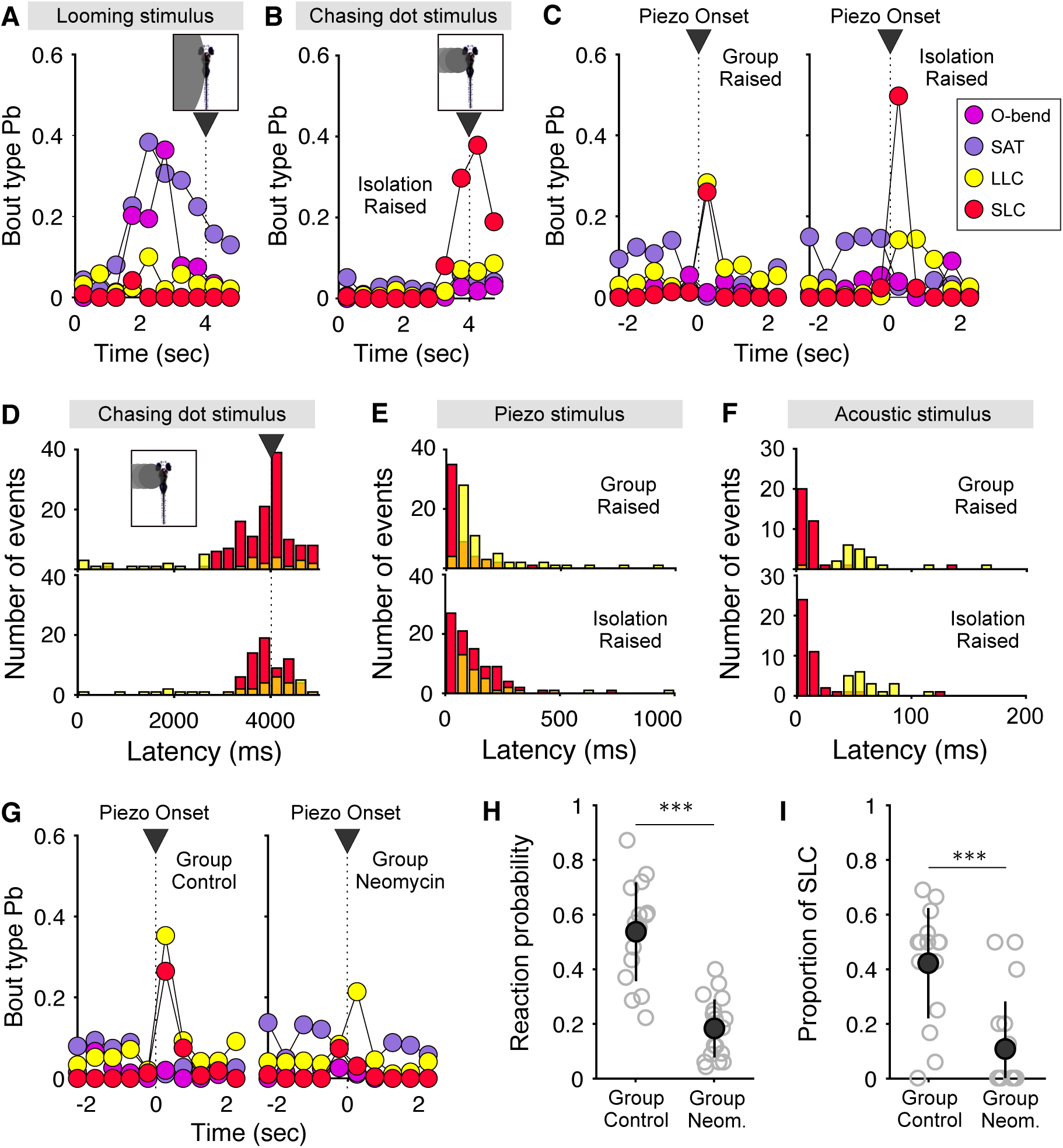
Response comparison between controlled visual and lateral line stimuli. (A,B) Probability of performing an avoidance bout type out of all 13 bout types was calculated in 500 ms time windows during presentation of a visual stimulus, which reaches the centroid of the larvae after 4 sec. Shown is the data of isolation raised larvae; N = 17 for looming stimuli (A) and N = 15 for chasing dot stimuli (B). Bout types are color-coded as shown in the legend. SAT: Shadow avoidance turn; LLC: Long latency C-start; SLC: Short latency C-start. (C) Bout type probability before and after presentation of local water vibrations through a semi-submerged rod, attached to a piezo bender (40V). Shown are the mean values of group (left, N = 15) and isolation raised larvae (right, N = 15). (D-F) Histograms of C-start latency after stimulus presentation shown for group (top) and isolation raised larvae (bottom). SLCs and LLCs are color-coded in red and yellow, respectively. The acoustic stimulus (F) consisted of a 100ms long 600 Hz pure tone played through speakers directly attached to the walls of the swimming arena. (G) Avoidance bout type probability across time before and after local water vibrations of 80V piezo input voltage is shown for control group raised larvae (left) and group raised larvae treated with neomycin to ablate the lateral line (right). (H) Reaction probability is defined as the number of SLC or LLC performed per total number of chasing dot stimulus presentations per larvae or either raising condition. (I) The proportion of SLCs was calculated as the number of stimulus induced SLCs divided by the sum over stimulus-induced SLCs and LLCs. Light gray open circles show every replicate per condition, dark gray filled circles and error-bars signal mean ± SD. Asterisk specify the results of two-tailed unsigned Wilcoxon tests between raising conditions (alpha level 0.05).

### Tables with statistical results

**Table S1:** Statistical comparison of group and isolation-raised larvae in their social avoidance measures. Figure reference indicates where the data is displayed. Avoidance area, C-Starts per bout and SLC proportion were calculated as described in the methods. Two-tailed unsigned Wilcoxon tests were applied for determining z and p values. MD and CI refer to the mean difference and its confidence intervals of the difference between group and isolation-raised larvae, as determined by 1000 bootstraps using the measure of effect size toolbox in Matlab. *Note that with the comparison of avoidance area measure in groups of seven larvae it was not possible to calculate z-values due to insufficient sample sizes.

| FigureRef | Measure | zValue | p | MD | $CI_{low}$ | $CI_{high}$ | $N_{gr}$ | $N_{iso}$ |
| --- | --- | --- | --- | --- | --- | --- | --- | --- |
| Fig. 1C | Avoidance area | NaN* | 0.00078 | -40.93 | -55.8 | -25.3 | 9 | 9 |
| Fig. S1C | Avoidance area | -2.7 | 0.0069 | -0.24 | -0.39 | -0.08 |  |  |
| Fig. 2E | C-starts per bout | -1.09 | 0.2765 | 0.21 | 0.19 | 0.24 | 43 | 43 |
| Fig. 2F | SLC proportion | -5.63 | <0.0001 | -0.17 | -0.22 | -0.13 |  |  |

**Table S2:** Statistical comparison for data shown in Fig 2B. Statistical comparison per bout type of the difference between group and isolation-raised larvae in their median inter-fish distance per larval test pair. N = 43 for both group and isolation-raised data. Two-tailed unsigned Wilcoxon tests were applied for each bout type for determining z and p values. To correct for multiple comparison, p-values were corrected (corr p) for the number of comparisons (13) using Holm’s sequential Bonferroni procedure. Mean and standard deviation (SD) are reported for each measure used for the comparison. AS: Approach Swim; Long CS: Long capture swim; BS: Burst swim; HAT: High angle turn; RT: Routine turn; SAT: Shadow avoidance turn; LLC: Long latency C-start; SLC: Short latency C-start.

| **Bout Type** | $Mean_{Gr}$ | $SD_{Gr}$ | $Mean_{Iso}$ | $SD_{Iso}$ | **zValue** | **p** | **corr p** |
| --- | --- | --- | --- | --- | --- | --- | --- |
| AS | 10.983 | 1.959 | 11.175 | 1.24 | -0.15 | 0.8833 | 1.1256 |
| Slow 1 | 11.673 | 0.827 | 11.274 | 1.15 | 1.58 | 0.114 | 0.5699 |
| Slow 2 | 11.821 | 0.778 | 11.231 | 1.328 | 2.73 | 0.0063 | 0.0571 |
| Short CS | 12.088 | 0.864 | 11.172 | 1.416 | 3.53 | 0.0004 | 0.0049 |
| J turn | 11.78 | 0.742 | 11.336 | 1.176 | 1.97 | 0.0489 | 0.3425 |
| HAT | 11.689 | 0.71 | 11.44 | 1.037 | 1.2 | 0.2299 | 0.9197 |
| Routine turn | 11.723 | 0.732 | 11.278 | 1.205 | 2.13 | 0.0329 | 0.2632 |
| SAT | 11.521 | 1.226 | 11.914 | 1.31 | -0.92 | 0.3599 | 1.0798 |
| Burst swim | 7.62 | 2.109 | 7.922 | 2.184 | -0.58 | 0.5628 | 1.1256 |
| Long CS | 5.991 | 3.249 | 7.384 | 3.899 | -1.72 | 0.0857 | 0.5139 |
| O bend | 6.618 | 2.361 | 8.246 | 2.438 | -2.87 | 0.0041 | 0.0455 |
| LLC | 6.998 | 1.983 | 8.177 | 2.117 | -2.82 | 0.0049 | 0.0487 |
| SLC | 3.788 | 1.142 | 4.978 | 1.798 | -4.1 | 0 | 0.0005 |

**Table S3:** Statistical comparison for data shown in Fig 2D. Statistical comparison per bout type of the difference between group and isolation-raised larvae in their probability of usage when the fish are at least within a distance of 10mm of one another. Bout type probability was determined as the number of the bouts per type divided by the total number of bouts swum of all 13 types. Two-tailed unsigned Wilcoxon tests were applied for each bout type for determining z and p values. To correct for multiple comparison, p-values were corrected (corr p) for the number of comparisons (5) using Holm’s sequential Bonferroni procedure. Mean and standard deviation (SD) are reported for each measure used for the comparison. N = 43 for both group raised and isolation raised data. Long CS: Long capture swim; LLC: Long latency C-start; SLC: Short latency C-start.

| **Bout Type** | $Mean_{Gr}$ | $SD_{Gr}$ | $Mean_{Iso}$ | $SD_{Iso}$ | **zValue** | **p** | **corr p** |
| --- | --- | --- | --- | --- | --- | --- | --- |
| Long CS | 0.015 | 0.006 | 0.016 | 0.024 | 2.36 | 0.0184 | 0.0551 |
| Burst swim | 0.023 | 0.016 | 0.024 | 0.026 | 0.5 | 0.6164 | 0.6164 |
| O bend | 0.014 | 0.009 | 0.019 | 0.011 | -2.36 | 0.0184 | 0.0551 |
| LLC | 0.071 | 0.03 | 0.051 | 0.025 | 4.02 | 0.0001 | 0.0003 |
| SLC | 0.036 | 0.014 | 0.064 | 0.037 | -3.84 | 0.0001 | 0.0005 |

**Table S4:**
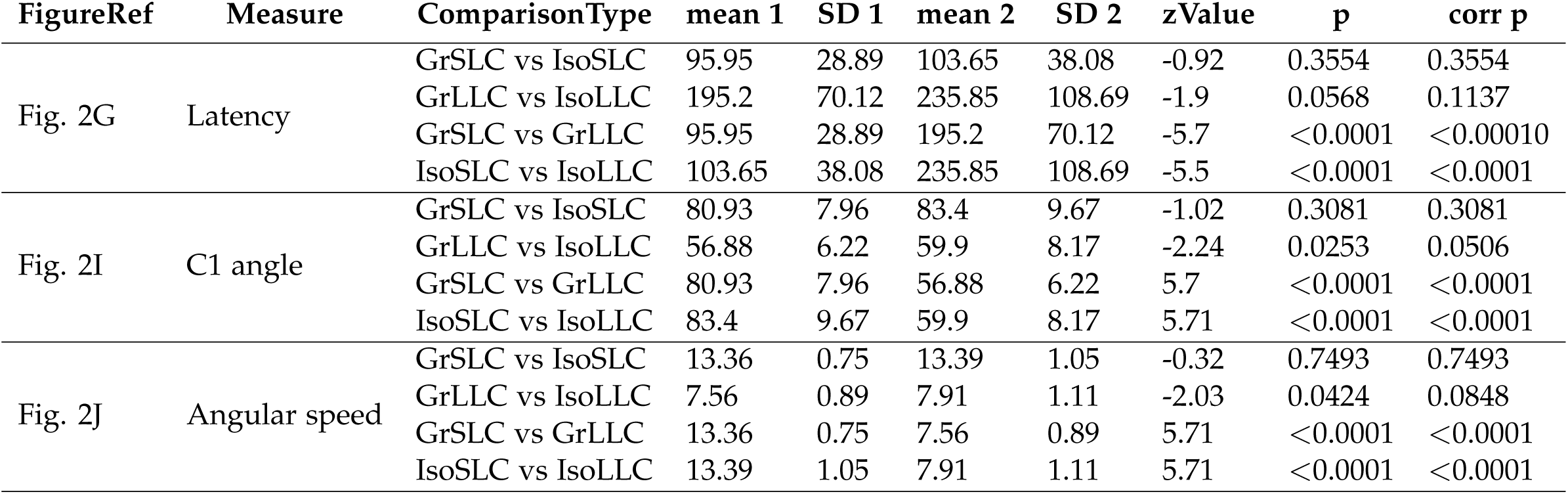
Statistical comparison of C-start types between group and isolation-raised larvae. Figure reference indicates where the data is displayed. Two-tailed unsigned and signed Wilcoxon tests were applied when comparing between raising conditions and between bout types within raising condition (repeated measure), respectively, for determining z and p values. To correct for multiple comparison, p-values were corrected (corr p) for the number of comparisons (4) using Holm’s sequential Bonferroni procedure. The comparison type is abbreviated as the following; GrLLC and GrSLC: Long and short latency C-starts performed by group-raised larvae; IsoLLC and IsoSLC: Long and short latency C-starts performed by isolation-raised larvae. Mean 1 refers to the first mentioned measure (i.e. GrLLC when written: GrLLC vs GrSLC) and likewise, mean 2 refers to the second mentioned measure. Similarly, SD 1 and 2 refer to the standard deviation of the measures.

**Table S5:** Statistical comparison for data shown in Fig S2 A comparing raising conditions for social and non-social testing conditions. Statistical comparison of difference between group and isolation raised larvae in the overall bout type Pb of all 13 bout types. N = 20 for both group raised and isolation raised data, tested in pairs or as single larvae. Two-tailed unsigned Wilcoxon tests were applied between raising conditions for each bout type for determining z and p values. To correct for multiple comparison, p-values were corrected (corr p) for the number of comparisons (13) using Holm’s sequential Bonferroni procedure. Mean and standard deviation (SD) are reported for each measure used for the comparison. AS: Approach Swim; Long CS: Long capture swim; BS: Burst swim; HAT: High angle turn; RT: Routine turn; SAT: Shadow avoidance turn; LLC: Long latency C-start; SLC: Short latency C-start.

| Bout Type | $Mean_{Gr}$ | $SD_{Gr}$ | $Mean_{Iso}$ | $SD_{Iso}$ | zValue | p | corr p |
| --- | --- | --- | --- | --- | --- | --- | --- |
| Tested in pairs of larvae |  |  |  |  |  |  |  |
| AS | 0.022 | 0.026 | 0.052 | 0.056 | -1.69 | 0.0909 | 0.5151 |
| Slow 1 | 0.273 | 0.083 | 0.261 | 0.088 | 0.09 | 0.9246 | 1.1957 |
| Slow 2 | 0.265 | 0.075 | 0.206 | 0.089 | 2.34 | 0.0193 | 0.1543 |
| Short CS | 0.023 | 0.01 | 0.019 | 0.009 | 1.58 | 0.1136 | 0.4545 |
| J turn | 0.053 | 0.012 | 0.065 | 0.026 | -1.72 | 0.0859 | 0.5151 |
| HAT | 0.122 | 0.022 | 0.11 | 0.039 | 1.47 | 0.1404 | 0.4542 |
| Routine turn | 0.19 | 0.068 | 0.173 | 0.059 | 0.53 | 0.5979 | 1.1957 |
| SAT | 0.023 | 0.018 | 0.045 | 0.025 | -3.29 | 0.001 | 0.0101 |
| Burst swim | 0.003 | 0.003 | 0.013 | 0.013 | -3.99 | 0.0001 | 0.0007 |
| Long CS | 0.002 | 0.002 | 0.005 | 0.004 | -2.02 | 0.0439 | 0.3072 |
| O bend | 0.003 | 0.002 | 0.009 | 0.004 | -4.34 | <0.0001 | 0.0002 |
| LLC | 0.014 | 0.015 | 0.025 | 0.016 | -3.18 | 0.0015 | 0.0133 |
| SLC | 0.006 | 0.003 | 0.017 | 0.011 | -4.23 | <0.0001 | 0.0003 |
| Tested as single larva |  |  |  |  |  |  |  |
| AS | 0.024 | 0.031 | 0.057 | 0.086 | -0.58 | 0.5609 | 2.03 |
| Slow 1 | 0.249 | 0.122 | 0.236 | 0.136 | 0.58 | 0.5609 | 1.6826 |
| Slow 2 | 0.303 | 0.128 | 0.244 | 0.151 | 1.04 | 0.2977 | 2.3814 |
| Short CS | 0.021 | 0.011 | 0.014 | 0.011 | 2.23 | 0.0256 | 0.3077 |
| J turn | 0.057 | 0.03 | 0.066 | 0.085 | 0.8 | 0.4249 | 2.9742 |
| HAT | 0.132 | 0.06 | 0.119 | 0.073 | 0.66 | 0.5075 | 2.4083 |
| Routine turn | 0.178 | 0.076 | 0.184 | 0.098 | -0.04 | 0.9676 | 1.1217 |
| SAT | 0.024 | 0.04 | 0.046 | 0.046 | -2.93 | 0.0033 | 0.0434 |
| Burst swim | 0.001 | 0.001 | 0.001 | 0.002 | 2.04 | 0.0416 | 0.4575 |
| Long CS | 0.003 | 0.005 | 0.003 | 0.006 | 0.7 | 0.4817 | 2.6071 |
| O bend | 0.002 | 0.002 | 0.004 | 0.006 | -1.37 | 0.1717 | 1.5454 |
| LLC | 0.006 | 0.008 | 0.025 | 0.055 | -1.56 | 0.1198 | 1.1984 |
| SLC | 0.001 | 0.001 | 0.001 | 0.002 | 0.78 | 0.4345 | 2.9742 |

**Table S6:** Statistical comparison for data shown in Fig S2 B-G and J-K. Figure reference indicates where the data is displayed. Total number of bouts swum and average bout duration were compared between raising conditions or between control and apomorphin treatment (Apo). Two-tailed unsigned Wilcoxon tests were applied for determining z and p values. MD and CI refer to the mean difference and its confidence intervals of the difference between group and isolation-raised larvae, as determined by 1000 bootstraps using the measure of effect size toolbox in Matlab.

| FigureRef | Treatment | Measure | zValue | p | MD | CI<sub>low</sub> | CI<sub>high</sub> | N |
| --- | --- | --- | --- | --- | --- | --- | --- | --- |
| Fig. S2B | Pairs | Bouts swum | 4.1 | <0.0001 | 24.78 | 15.43 | 32.91 | Gr: 20 |
| Fig. S2C |  | Bout dur. | -3.02 | 0.0026 | -13.35 | -21.17 | -6.15 | Iso: 20 |
| Fig. S2D | Alone | Bouts swum | 3.18 | 0.0015 | 25.41 | 11.89 | 38.75 | Gr: 20 |
| Fig. S2E |  | Bout dur. | -2.75 | 0.006 | -17.52 | -34.98 | -0.13 | Iso: 20 |
| Fig. S2F | Apo. | Bouts swum | 4.1 | <0.0001 | 24.78 | 15.66 | 33.6 | Ctr: 20 |
| Fig. S2G |  | Bout dur. | -3.02 | 0.0026 | -13.35 | -20.62 | -5.76 | Apo: 20 |
| Fig. S2J | Iso3 | Bouts swum | 6.55 | <0.0001 | 26.22 | 20.69 | 32.14 | Gr: 40 |
| Fig. S2K |  | Bout dur. | -0.31 | 0.7574 | -1.89 | -6.11 | 2.23 | Iso: 39 |

**Table S7:**
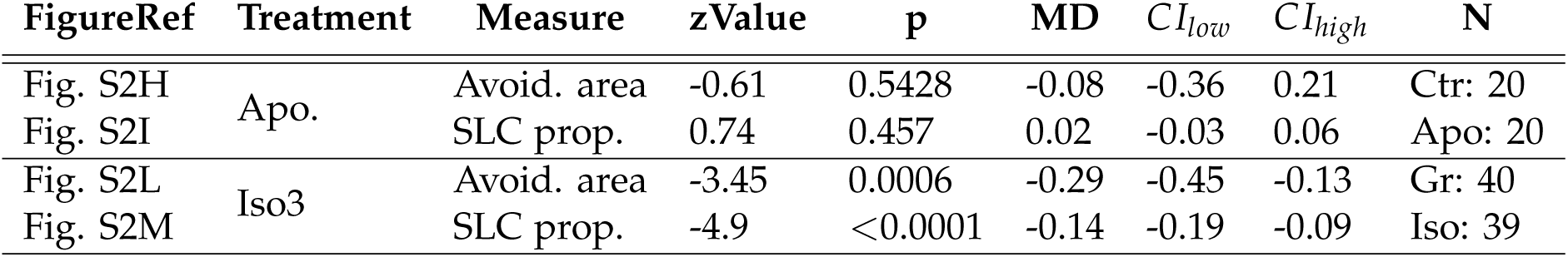
Statistical comparison for data shown in Fig S2 H-I and L-M. Social avoidance measures after apomorphin treatment (Apo.) or larvae that were isolated from 3dpf until testing (Iso3) were compared to regular group raised control larvae. Figure reference indicates where the data is displayed. Two-tailed unsigned Wilcoxon tests were applied for determining z and p values. MD and CI refer to the mean difference and its confidence intervals of the difference between group and isolation-raised larvae, as determined by 1000 bootstraps using the measure of effect size toolbox in Matlab.

**Table S8:** Statistical comparison of angular distributions of the C-start preceding bout of the non-focal larvae. Figure reference indicates where the data is displayed. Two-tailed unsigned and signed Wilcoxon tests were applied when comparing between raising conditions and between bout types within raising condition (repeated measure), respectively, for determining z and p values. To correct for multiple comparison, p-values were corrected (corr p) for the number of comparisons (4) using Holm’s sequential Bonferroni procedure. The comparison type is abbreviated as the following; GrLLC and GrSLC: Long and short latency C-starts performed by group-raised larvae; IsoLLC and IsoSLC: Long and short latency C-starts performed by isolation-raised larvae. Mean 1 refers to the first mentioned measure (i.e. GrLLC when written: GrLLC vs GrSLC) and likewise, mean 2 refers to the second mentioned measure. Similarly, SD 1 and 2 refer to the standard deviation of the measures. Matlab’s CircStat toolbox was used to calculate Kuiper k.

| FigureRef | Comparison Type | median 1 | SD 1 | median 2 | SD 2 | Kuiper k | p | corr p |
| --- | --- | --- | --- | --- | --- | --- | --- | --- |
| Fig. 3A | GrSLC vs IsoSLC | 0.83 | 0.86 | 1.22 | 0.88 | 200627 | 0.001 | 0.004 |
|  | GrLLC vs IsoLLC | 1.26 | 0.91 | 1.26 | 0.87 | 304129 | 0.005 | 0.01 |
|  | GrSLC vs GrLLC | 0.83 | 0.86 | 1.26 | 0.91 | 292750 | 0.001 | 0.004 |
|  | IsoSLC vs IsoLLC | 1.22 | 0.88 | 1.26 | 0.87 | 82499 | >0.1 | >0.1 |
| Fig. 3D | GrSLC vs IsoSLC | 0.9 | 0.89 | 1.01 | 0.86 | 18555 | >0.1 | >0.1 |
|  | GrLLC vs IsoLLC | 0.95 | 0.86 | 0.9 | 0.82 | 54285 | >0.1 | >0.1 |
|  | GrSLC vs GrLLC | 0.9 | 0.89 | 0.95 | 0.86 | 41320 | >0.1 | >0.1 |
|  | IsoSLC v IsoLLC | 1.01 | 0.86 | 0.9 | 0.82 | 23052 | >0.1 | >0.1 |
| Fig. 3G | GrSLC vs IsoSLC | 1.22 | 0.84 | 1.16 | 0.79 | 3719 | >0.1 | >0.1 |
|  | GrLLC vs IsoLLC | 1.32 | 0.81 | 1.33 | 0.71 | 26768 | 0.05 | >0.1 |
|  | GrSLC vs GrLLC | 1.22 | 0.84 | 1.32 | 0.81 | 16315 | >0.1 | >0.1 |
|  | IsoSLC vs IsoLLC | 1.16 | 0.79 | 1.33 | 0.71 | 6012 | >0.1 | >0.1 |
| Fig. S3A | GrSLC vs IsoSLC | 1.47 | 0.84 | 1.27 | 0.79 | 7760 | >0.1 | >0.1 |
|  | GrLLC vs IsoLLC | 1.3 | 0.79 | 1.18 | 0.77 | 19604 | >0.1 | >0.1 |
|  | GrSLC vs GrLLC | 1.47 | 0.84 | 1.3 | 0.79 | 15644 | >0.1 | >0.1 |
|  | IsoSLC vs IsoLLC | 1.27 | 0.79 | 1.18 | 0.77 | 11990 | >0.1 | >0.1 |
| Fig. S3D | GrSLC vs IsoSLC | 1.07 | 0.89 | 1.13 | 0.89 | 36543 | >0.1 | >0.1 |
|  | GrLLC vs IsoLLC | 1.17 | 0.89 | 1.14 | 0.86 | 63758 | >0.1 | >0.1 |
|  | GrSLC vs GrLLC | 1.07 | 0.89 | 1.17 | 0.89 | 72195 | >0.1 | >0.1 |
|  | IsoSLC vs IsoLLC | 1.13 | 0.89 | 1.14 | 0.86 | 33006 | >0.1 | >0.1 |

**Table S9:** Statistical comparison for data shown in Fig 3 and S3. Social avoidance measures were compared between raising conditions after manipulation of the sensory input, i.e. Neomycin (Neom.) and *CuSO*_4_ treatment for ablation of the LL. Figure reference indicates where the data is displayed. Two-tailed unsigned Wilcoxon tests were applied for determining z and p values. MD and CI refer to the mean difference and its confidence intervals of the difference between group and isolation-raised larvae, as determined by 1000 bootstraps using the measure of effect size toolbox in Matlab.

| FigureRef | Treatment | Measure | zValue | p | MD | CI<sub>low</sub> | CI<sub>high</sub> | N |
| --- | --- | --- | --- | --- | --- | --- | --- | --- |
| Fig. 3B | Contr. | Avoid. area | -2.61 | 0.009 | -0.28 | -0.47 | -0.08 | Gr: 31 |
| Fig. 3C |  | SLC prop. | -5.66 | <0.0001 | -0.19 | -0.24 | -0.14 | Iso: 39 |
| Fig. 3E | Dark | Avoid. area | -1.19 | 0.2353 | -0.06 | -0.2 | 0.09 | Gr: 27 |
| Fig. 3F |  | SLC prop. | -4.82 | <0.0001 | -0.18 | -0.23 | -0.12 | Iso: 28 |
| Fig. 3H | Neom. | Avoid. area | -1.08 | 0.2807 | -0.12 | -0.33 | 0.09 | Gr: 19 |
| Fig. 3I |  | SLC prop. | -3.02 | 0.0025 | -0.11 | -0.18 | -0.04 | Iso: 18 |
| Fig. S3B | CuSO<sub>4</sub> | Avoid. area | -0.6 | 0.5501 | -0.07 | -0.28 | 0.14 | Gr: 27 |
| Fig. S3C |  | SLC prop. | 0.39 | 0.6986 | 0 | -0.07 | 0.07 | Iso: 28 |
| Fig. S3E | Handled | Avoid. area | -1.99 | 0.0468 | -0.27 | -0.5 | -0.03 | Gr: 20 |
| Fig. S3F |  | SLC prop. | -3.39 | 0.0007 | -0.14 | -0.21 | -0.07 | Iso: 20 |

**Table S10:** Statistical comparison for data shown in Fig. 4 and Fig. S4E and F. Difference between group and isolation raised larvae in their escape bout reactions to looming stimuli, an approaching dark spot (CD) and local water vibrations caused by a semi-submerged capillary tip attached to a piezo bender (piezo). Piezo neom. refers to the comparison between control fish and neomycin treated group raised larvae for ablation of the LL. Figure reference indicates where the data is displayed. Two-tailed unsigned Wilcoxon tests were applied for determining z and p values. MD and CI refer to the mean difference and its confidence intervals of the difference between group and isolation-raised larvae, as determined by 1000 bootstraps using the measure of effect size toolbox in Matlab.

| FigureRef | Stim type | Measure | zValue | p | MD | CI<sub>low</sub> | CI<sub>high</sub> | N |
| --- | --- | --- | --- | --- | --- | --- | --- | --- |
| Fig. 4B | Loom | ReactionPb | 0.79 | 0.4278 | 0.030 | -0.020 | 0.08 | Gr: 16 |
| Fig. 4C |  | O-bend Prop. | 0.9 | 0.3678 | 0.06 | -0.04 | 0.14 | Iso: 17 |
| Fig. 4E | CD | ReactionPb | 2.96 | 0.0031 | 0.20 | 0.10 | 0.32 | Gr: 14 |
| Fig. 4F |  | SLC Prop. | 1.18 | 0.2384 | 0.11 | -0.06 | 0.28 | Iso: 15 |
| Fig. 4H | Piezo | ReactionPb | -1.14 | 0.254 | -0.090 | -0.23 | 0.07 | Gr: 15 |
| Fig. 4I |  | SLC Prop. | -3.15 | 0.0016 | -0.34 | -0.48 | -0.18 | Iso: 15 |
| Fig. S4H | Piezo | ReactionPb | 4.36 | <0.0001 | 0.36 | 0.24 | 0.46 | Ctr: 15 |
| Fig. S4I | Neom. | SLC Prop. | 3.73 | 0.0002 | 0.32 | 0.18 | 0.44 | Neom: 18 |

**Table S11:**
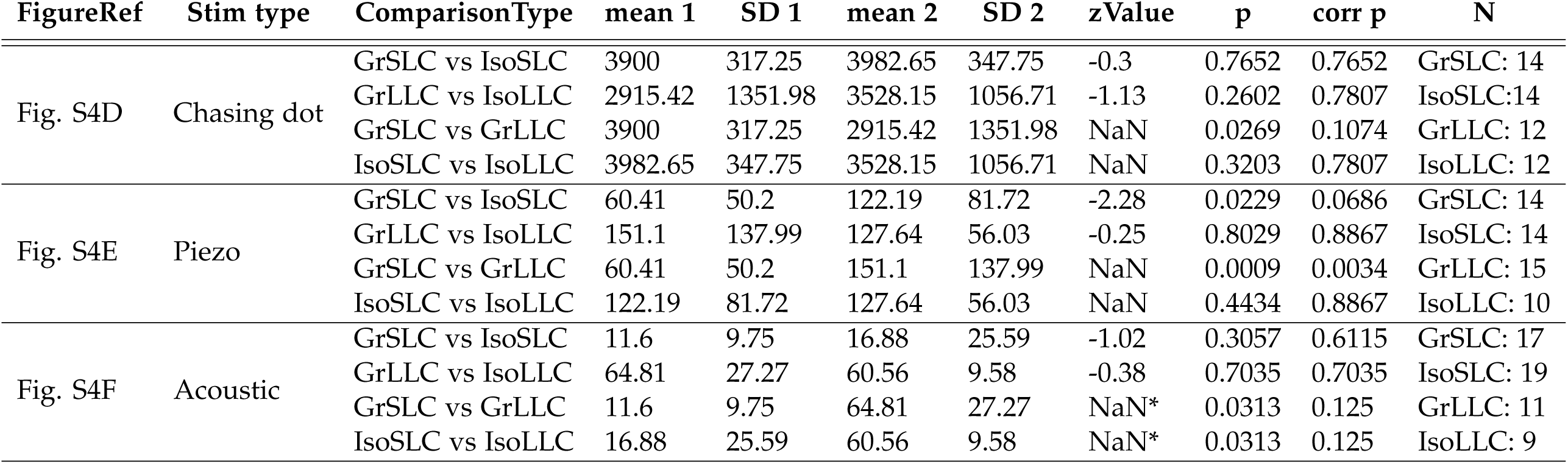
Comparison of C-start latency after stimulus presentation shown in Fig. S4 DF. Figure reference indicates where the data is displayed. Two-tailed unsigned and signed Wilcoxon tests were applied when comparing between raising conditions and between bout types within raising condition (repeated measure), respectively, for determining z and p values. To correct for multiple comparison, p-values were corrected (corr p) for the number of comparisons (4) using Holm’s sequential Bonferroni procedure. The comparison type is abbreviated as the following; GrLLC and GrSLC: Long and short latency C-starts performed by group-raised larvae; IsoLLC and IsoSLC: Long and short latency C-starts performed by isolation-raised larvae. Mean 1 refers to the first mentioned measure (i.e. GrLLC when written: GrLLC vs GrSLC) and likewise, mean 2 refers to the second mentioned measure. Similarly, SD 1 and 2 refer to the standard deviation of the measures. *Note that for the repeated measure comparison between C-start bout types within raising condition it was not possible to calculate z-values due to insufficient sample sizes. The sample sizes may differ from the total number of animals tested, since only animals who performed a C-start during the stimulus presentation could be taken into consideration, see Fig. 4 E-F and H-I.

